# Fragment-Based Drug Discovery for Transthyretin Kinetic Stabilisers Using a Novel Capillary Zone Electrophoresis Method

**DOI:** 10.1101/2025.02.22.639275

**Authors:** Wenjie Chen

**Author notes:** Current Address: Research Department of Pharmaceutical & Biological Chemistry, School of Pharmacy, University College London, 29-39 Brunswick Square, London WC1N 1AX.

## Abstract

A Capillary Zone Electrophoresis (CZE) fragment screening methodology was developed and applied to the human plasma protein Transthyretin (TTR), normally soluble, but could misfold and aggregate, causing amyloidosis. Termed Free Probe Peak Height Restoration (FPPHR), it monitors changes in the level of free ligand known to bind TTR (the Probe Ligand) in the presence of competing fragments. 129 fragments were screened, 12 of the 16 initial hits (12.4% hit rate) were co-crystallised with TTR, 11 were found at the binding site (92% confirmation rate). Subsequent analogue screens have identified a novel TTR-binding scaffold 4-(3H-pyrazol-4-yl)quinoline and its derived compounds were further studied by crystallography, Circular Dichroism (CD), Isothermal Titration Calorimetry (ITC) and radiolabelled ^125^I-Thyroxine displacement assay in neat plasma. Two lead molecules had similar ITC K_d_ and ^125^I-Thyroxine displacement IC_50_ values to that of Tafamidis, adding another potential pipeline for transthyretin amyloidosis. The methodology is reproducible, procedurally simple, automatable, label-free without target immobilisation, non-fluorescence based and site-specific with low false positive rate, which could be applicable to fragment screening of many drug targets.

## Introduction

With seven approved fragment-derived drugs and many others currently in clinical trials [1], Fragment-Based Drug Discovery (FBDD) is a widely accepted strategy alongside High-Throughput Screening (HTS) in the field of early-stage drug discovery. As well as the need of a reliable system for obtaining ligand-protein structural data, another pre-requisite for the success of FBDD is the ability to detect weakly binding fragments. There is a range of weak molecular interaction detection techniques for fragment screening, including but not limited to NMR [2, 3], X-ray crystallography [4], Microscale Thermophoresis (MST) [5], Mass Spectrometry [6], and Surface Plasmon Resonance (SPR) [7], which is the mainstream technique used in the biopharma sector. Each technique holds a unique set of advantages and disadvantages, thus there is no one-technique-for-all and meticulous considerations should be given when choosing, based on target biochemical properties and overall objective of the screening campaign.

Capillary Electrophoresis (CE) being both biophysical and chromatographic was first documented in 1930 [8] and has since formed a new branch in separation science. It is now a well-established analytical technique mostly used in diagnostic and clinical settings such as haemoglobinopathy investigations, DNA genotyping and pharmaceutical drug analysis. The CE apparatus [9] is simple, consisting of a thin glass tube (the capillary), voltage supply, electrolyte buffer reservoirs, and a detector (**Fig 1**). There is also a variety of separation modes in CE adapted to separate a wide range of charged and neutral analytes including inorganic ions, small molecules, peptides, nucleic acids and proteins. For example, Isoelectric Focusing, Isotachophoresis, Capillary Zone Electrophoresis (CZE), Capillary Gel Electrophoresis (CGE) and Micellar Electrokinetic Chromatography, are all separation processes that take place in a thin glass tube but by fundamentally different mechanisms. Superior separation efficiency and reproducibility (**S1 Fig**), microscale sample consumption, low running costs, broad analyte applicability and automatability has made CE becoming more popular in the 1980s and most notable on sequencing of the human genome by CGE in the late 1990s.

**Fig 1.**
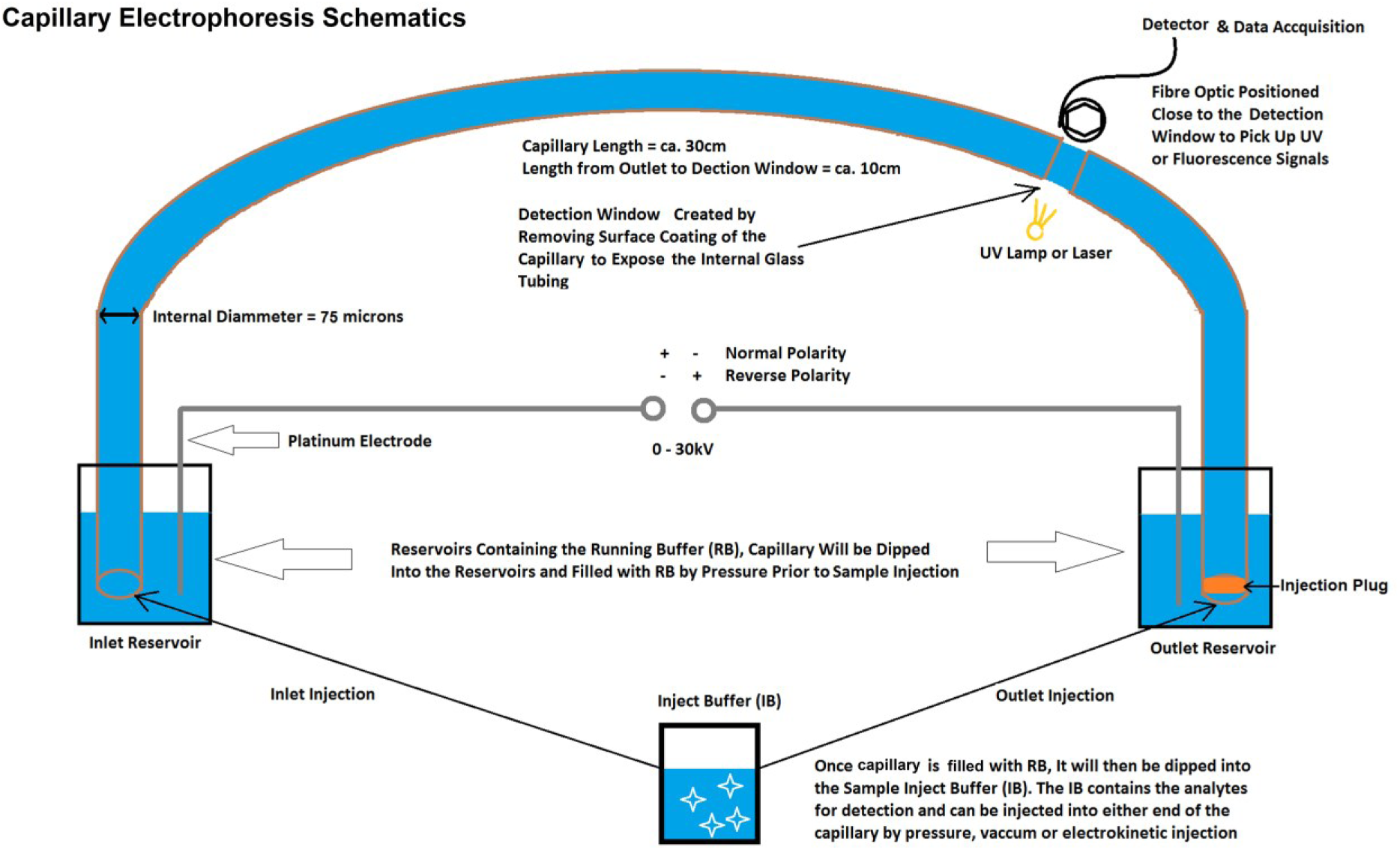
Schematic Representation of a Typical CE Apparatus

Unlike conventional chromatography, CZE separates analytes in a homogenous buffer and fully solution-based environment without a stationary phase, providing the framework for Affinity Capillary Electrophoresis (ACE). Its specialised use in fragment screening was initially practiced by the biotech company Cetek Corporation in Massachusetts, US in around 2005. A published example was a CZE fragment screening assay for Heat Shock Protein 90α ATPase domain [10] developed based on an ACE method called CEfrag™ [11]. The assay detects fragment-protein binding by monitoring the mobility shift of the Probe Ligand, Radicicol, a natural antibiotic known to bind Hsp90. Various binding and/or screening ACE assays [12–16] for multiple targets across different disease areas has since been developed showcasing the immense versatility in CE assay method development. As a novel variant to ACE methods previously described, here we present an uncomplicated and automatable fragment screening method for **Trans**port Protein of **Thy**roxine and **Retin**ol - Transthyretin.

Transthyretin (TTR) is a homotetrameric plasma protein (also present in cerebrospinal fluid) with two thyroxine binding sites [17] (**S2 Fig**). Thyroxine in the plasma is however predominantly bound by Thyroxine Binding Globulin and albumin, the transthyretin binding sites are mostly empty [18]. In the absence of bound ligand, the tetramer is prone to dissociation and subunit misfolding that generates the precursors [19] for amyloid fibre formation. These cross-β sheet structured fibres [20] are deposited in tissues, commonly in the peripheral nervous system and the heart, which leads to detrimental neurological and cardiac complications [21]. Some individuals are especially susceptible to TTR amyloidosis because they inherited mutations [22] in the transthyretin gene and the expressed TTR variants are less stable. Old age can also cause TTR amyloidosis of which wild type TTR fibres are found in the heart [23]. The prospects for developing small molecules as kinetic stabilisers of the TTR tetramer and thus as amyloidosis inhibitors was recognised more than twenty years ago with structure-based drug design ventures published [24, 25]. This has led to the development of Tafamidis, the first approved small molecule drug for the treatment of Familial Amyloid Polyneuropathy in 2011. Continual drug discovery efforts for transthyretin amyloidosis were made, resulting recent approval of several TTR gene-silencing agents in 2022 and anti-TTR monoclonal antibodies as well as small molecule stabilisers currently in clinical trial [26].

The CZE TTR fragment screening assay described here employs 8-Anilino-1-Naphthalenesulfonic acid (8-ANS) as the Probe Ligand, which binds at the TTR thyroxine binding sites (PDB 3CFN [27], K_d1_ = 1.05 µM K_d2_ = 4.79 µM [28]) . It detects the increase in unbound 8-ANS UV peak height when test fragment in the capillary encounters the injected 8-ANS-TTR complex, hence the name Free Probe Peak Height Restoration (FPPHR) method. Under optimised conditions, it provides consistent unbound 8-ANS migration time, peak height and area. This allows sensitive detection of subtle changes in the binding events involving TTR, 8-ANS and the competitors. Furthermore, the detection of 8-ANS displacement from TTR makes the fragment screening assay site-specific, a feature not found in many other fragment screening techniques. In contrast to other ACE methods, filling of the capillary with protein solution does not occur in FPPHR, and the protein-containing solution is never mixed with the screened fragments, reducing protein consumption and simplifying sample handling. Primary screening of 129 rule-of-3 compliant fragments by the CZE FPPHR method has identified 16 initial hits. 11 of the 12 TTR fragment hits selected based on novelty and diversity were validated by X-ray protein crystallography with the added benefit of structural characterisation. Follow-on analogue screenings led to discovery of 4-(3H-pyrazol-4-yl)quinoline and its derivatives were further tested by crystallography, circular dichroism (CD), isothermal titration calorimetry (ITC) and ^125^I-thyroxine displacement assay in neat plasma.

In summary, the research impact here is in two parts. The discovery of novel, tractable small molecule leads with mid-nanomolar (10 – 100 nM) affinity, has paved the way for development of an effective TTR kinetic stabiliser. Additionally, by showcasing CZE FPPHR method in detection of site-specific weakly-binding fragments, in a yes-or-no manner, whilst affording remarkably low rate of false-positive, significant shift in the understanding of its utilisation in FBDD is instigated. Drug discovery scientists from both academia and industry could adopt and apply the FPPHR method as a primary screening tool in their FBDD endeavours expanding the repertoire of future CZE assays. The high quality and reliable fragment hits from CZE FPPHR will invariably expedite the drug discovery process by considerably mitigating the risk of false-positives and irrelevant binders.

## Materials and Methods

### Chemicals, Reagents & Protein

Tafamidis and Compound SEL101928 were synthesised by Selcia Ltd. (Ongar, Essex, UK). Thyroxine T_4_ and 8-Anilino-1-naphthalenesulfonic acid ammonium salt (8-ANS) were purchased from Sigma (Poole, Dorset, UK). All other tested compounds were either from the chemical store of Selcia Ltd., the Selcia Fragment Library (SFL) or purchased from ChemBridge Corporation (San Diego, CA, USA), or Sigma (St. Louis, MO, USA).

The Selcia Fragment Library contains around 1300 rule-of-3 compliant fragments with purity of > 95% (LC-MS & 1^H^-NMR) dissolved in Dimethyl Sulfoxide (DMSO) to 30 mM as stocks solutions. Aqueous solubility was measured to be >1 mM by measuring A_650nm_ (< 0.01) after 3 hours of shaking at 25°C in 35 mM HEPES pH 7.8 at 1 mM, final DMSO % = 3.3 volume to volume (v/v).

Human transthyretin (UniProt P02766) lyophilised from 0.02 M NH_4_HCO_3_ solution was purchased from SCIPAC (Sittingbourne, Kent, UK), Product Code P171-1 Lot.1802-20-1, purity > 96%. Dried TTR was weighed and dissolved with CZE Running Buffers A or B to give a final stock concentration of 100 μM tetramers.

### Capillary Electrophoresis

CZE assays development and fragment screening were performed on a Beckman P/ACE MDQ capillary electrophoresis system (Beckman Coulter Inc., Brea, CA) operated by 32 Karat™ software. All capillaries (Polymicro Technologies, Phoenix, AZ) used were internally coated with polyvinyl alcohol and of around 30 cm in length, 75 microns in internal diameter. Capillary temperature was maintained at 20 °C by recirculating fluorinated fluid coolant during electrophoresis. Sample trays and buffer trays temperature were maintained at 4°C and ambient respectively. For every electrophoretic run, the capillary was pressure rinsed and prefilled with electrophoresis buffer also known as background electrolyte (termed as Running Buffer hereafter) before injection. After each electrophoretic run, the capillary was rinsed with Running Buffer and H_2_O by pressure (20 psi for 1 minute). Three Running Buffers (RB), **A:** 20 mM Tris pH 7.5, 1 mM KCl, 1.62 mM NaCl; **B:** 20 mM HEPES pH 7.0, 2 mM CaCl_2_ and **C:** 20 mM HEPES pH 7.0, 2 mM CaCl_2_, 1% DMSO were used. All CE traces were analysed by 32Karat™ Software with a built-in automatic peak integration algorithm.

### TTR FPPHR CZE Assay Development and Fragment Screening

**Control 1** Inject Buffer (IB) containing 30 μM 8-ANS was prepared by mixing 1 μL of 3 mM 8-ANS DMSO stock with 99 μL of Running Buffer B in a 0.2 mL PCR tube. **Control 2** Inject Buffer was prepared by mixing 1μL of 3 mM 8-ANS DMSO stock with 20 μL of 100 μM tetrameric TTR stock in Running Buffer B and 79 μL of Running Buffer B. The final concentrations (f/c) were 8-ANS 30 μM, TTR 20 μM and 1% DMSO v/v.

The maximum Free Probe Peak Height (Control 1 Peak Height) was determined by Outlet Injection (see **Fig 1**) of Control 1 with an injection pressure of 0.5 psi for 5 seconds (equivalent to ≈ 20 – 30 nL). An identical injection of Control 2 provides the minimum Free Probe Peak Height (Control 2 Peak Height) enabling calculation of % of bound and unbound *i.e.* free 8-ANS. To determine the 8-ANS peak height in Control 1 and Control 2, the capillary was prefilled with Buffer C and electrophoresis carried out at 10 kV (Normal Polarity, **Fig 1**) with UV monitoring at 230 nm. For displacement analysis, ligand DMSO stock was diluted in Buffer B and used to fill the capillary as the Running Buffer. Electrophoresis commenced after injection of Control 2, thus the injected 8-ANS-TTR complex migrates inside a capillary filled with the test ligand. The observed 8-ANS peak height is defined as Sample Peak Height and the displaced 8-ANS can be calculated as:

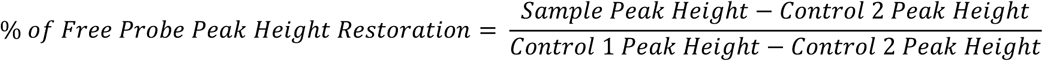

### Co-Crystallisation of TTR-Ligand Complex, Data Collection & Processing

Crystals were grown by vapor diffusion using a published crystallisation cocktail (100 mM HEPES pH 7.5, 200 mM CaCl_2_, 28% v/v PEG400) at 21 °C [29]. The hanging drop contained 1 μL of crystallisation cocktail, 1 μL of diluted ligand and 1 μL of TTR protein solution (85 μM). Final ligand concentration and DMSO % in crystal drop varied from 0.13 – 6.6 mM and 0.13 – 6.6 % respectively depending on their solubility and affinity. Crystals were harvested and flash cooled in a N_2_ cryostream or dipped into liquid ethane without cryoprotectant. Data was collected at Diamond Light Source (Oxon, UK) beamline I24, I04 or I04-1 (Pilatus 6M or ADSC Q315) or the European Synchrotron Radiation Facility (ESRF, Grenoble, France) beamline ID29 (Pilatus 6M-F) or in-house (Bruker X8 Diffractometer with APEXII CCD detector). Crystal unit cell was initially indexed by software MOSFLM [30] and LABELIT [31] within the EDNA [32] automatic data processing workflow. Diffraction image integration was performed by XDS [33] and reflection intensities were scaled by SCALA [34] or by PROTEUM2 if collected in-house. Phases were obtained by molecular replacement method using PHASER [35], 1DVQ from the Protein Data Bank (PDB) was used as the search model. Atomic positions of the initial model were refined by REFMAC5 [36]. Model building and ligand fitting was completed using COOT [37]. Model graphics were generated using CCP4MG [38]. Crystallographic statistics of all structures can be found in **S4 Table**.

### Induced Circular Dichroism (CD) of SFL001535

CD experiments were conducted in Diamond Light Source (Oxon, UK) beamline B23 installed with an Olis™ DSM 20 CD spectrophotometer. CD Buffer (10 mM Tris-HCl, 5 mM NaCl. pH 8.0) was prepared and loaded (450 μL) into a UV cuvette (10 mm path length). The near-UV (260 – 380 nm) CD spectrum of the solution was measured. 1 μL of SFL001535 10 mM DMSO stock was added into the cuvette and mixed by inversion. The CD spectrum of this solution was measured again. Contents of the cuvette containing SFL001535 in buffer were removed and the cuvette was washed with nitric acid. CD Buffer containing 13.62 μM of TTR was loaded into the cuvette and the near-UV CD spectrum of TTR was obtained (measured 10 times). A total of 8 μL of SFL001535 10 mM stock was added to the solution in 4 step additions (1 + 1 + 2 + 4 μL) and the CD spectrum were measured after each addition. The induced CD of SFL001535 was calculated by subtracting the CD spectrum of TTR alone.

### Isothermal Titration Calorimetry

All ITC experiments were conducted on an iTC200 calorimeter (GE Healthcare), and the settings were 25 °C, stir at 1000 RPM, 1 – 2 μL injection spaced with 100 – 150 seconds, reference power: 6 μcal/sec. Isotherms were fitted using ORIGIN 7 software (OriginLab, MA, USA). To prepare TTR ITC protein solution (5 mL), purified and freeze-dried human TTR was weighted and dissolved in 310 μL of pure water. The reconstituted protein solution was then desalted by spin column (Bio-Rad, Bio-Gel P6) and the eluted volume was transferred to a 15mL falcon tube. The protein solution was further diluted with 10x Phosphate Buffered Saline (PBS, 500 μL), DMSO (150 μL) and water, (f/c: 4.3 mM Na_2_HPO_4_, 137 mM NaCl, 2.7 mM KCl, 1.47 mM KH_2_PO_4_, 3% DMSO v/v, TTR 19.35 – 25 μM, pH 7.8). UV A260/280 was measured to obtain the actual protein concentration using a calculated extinction coefficient of 18450 M^-1^ cm^-1^. To prepare ligand ITC solution (1 mL), 30 μL of ligand stock in DMSO was diluted with 10x PBS (100 μL) and water (f/c: 4.3 mM Na_2_HPO_4_, 137 mM NaCl, 2.7 mM KCl, 1.47 mM KH_2_PO_4_, 3% DMSO v/v, Ligand 280 – 600 μM, pH 7.8).

## ^125^I-T_4_ Displacement Assay in Plasma

The assay was adapted from one described previously [39]. Briefly, test ligand and ^125^I-T_4_ (Perkin Elmer Inc., Waltham, MA) were mixed with undiluted normal human reference plasma (Precision Biologic Inc., Nova Scotia, Canada). The TTR in plasma was immunoprecipitated by sheep anti-human prealbumin polyclonal antibody (The Binding Site Group Ltd., Birmingham, UK). The immunoprecipitate was washed with buffer and their radioactivity measured with a 2470 WIZARD automatic gamma counter (Perkin Elmer Inc., Waltham, MA). Gamma counts of samples containing DMSO only and 714 μM of Tafamidis were used as the maximum and minimum respectively. Percentage of total radioligand binding for samples of different ligand concentration was then calculated as follows:

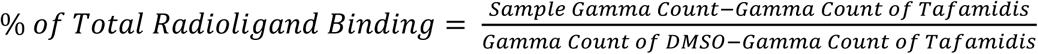

To calculate IC_50_ values, percentage of total radioligand binding at different concentration points were entered in SigmaPlot™ 11.0. Titration curves were done in triplicates with a minimum of 5 data points fitted by 4-Parameter Logistic Function. Representative graphs of 1 of the 3 fitted curves for each test ligand can be found in **S3 File**.

## Results

### Development of TTR CZE FPPHR Fragment Screening Assay: Effect of TTR Binding on Free 8-ANS UV Absorbance Peak Height and Area

The migration time, peak height and area of free 8-ANS (30 µM) were averaged around 2.3 minutes, 1640 and 10274 Arbitrary Units (AU) respectively with excellent replicability (**Fig 2**, **Trace A & B,**). When 8-ANS (30 μM) was mixed with TTR (20 μM) in a different vial of Inject Buffer and separated again, two peaks appeared (**Fig 2**, **Trace C & D**). The first peak corresponds to the migration time of free 8-ANS with a reduced average peak height and area of 665 and 6686 AU respectively. The second peak corresponding to the UV absorbance of TTR eluted at around 3.5 minutes was much broader and higher. Since the injection volume and concentrations were the same, the reduction in free (unbound) 8-ANS peak height and area was a result of molecular binding to TTR. The late eluting peak is the combined UV absorbance of apo TTR, singly and doubly 8-ANS bound TTR.

**Fig 2.**
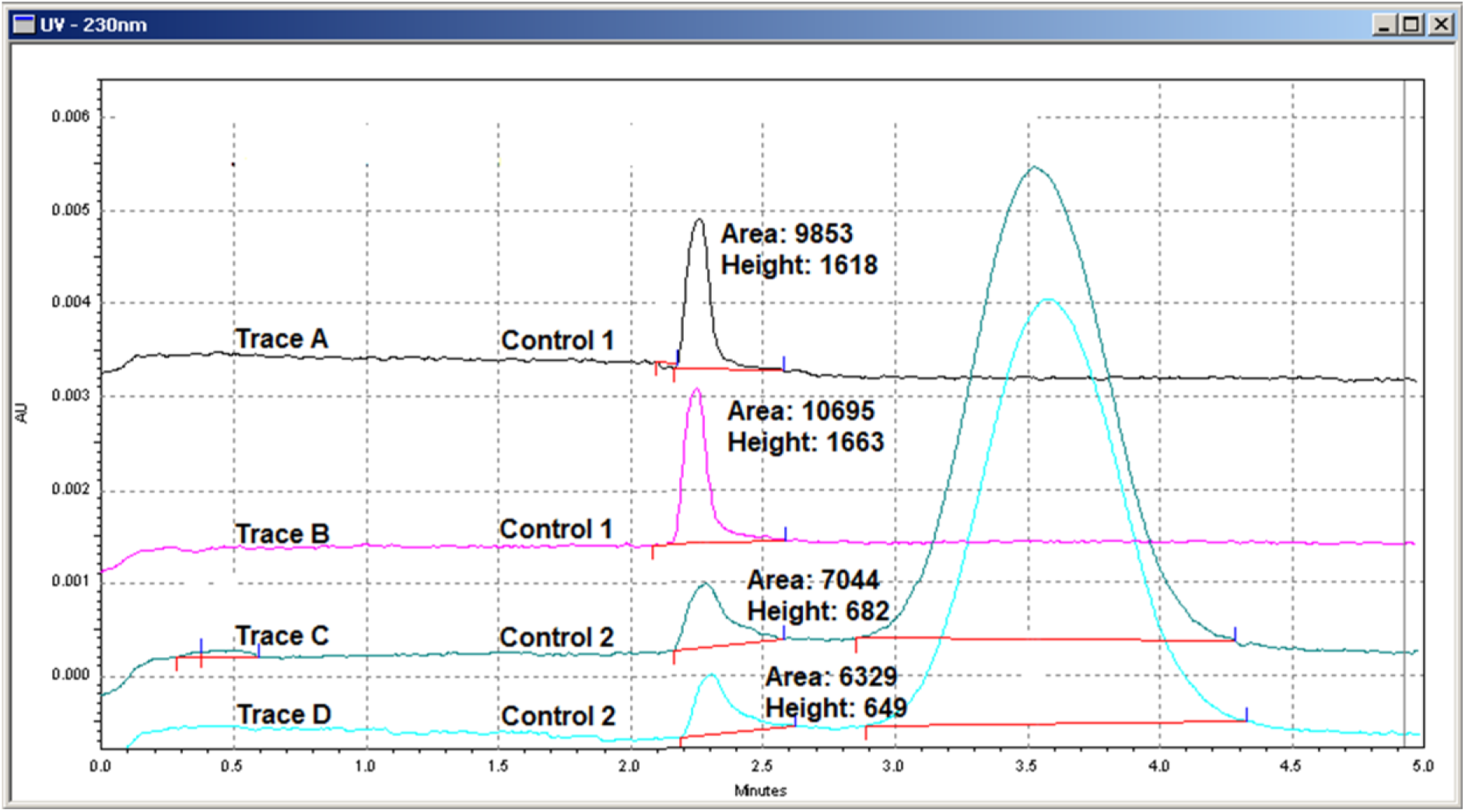
Effect of TTR Binding on Free 8-ANS UV Absorbance Peak Height and Area. Two vials of Inject Buffer were prepared, one contained 30 μM 8-ANS (Control 1) and the other contained 30 μM 8-ANS and 20 μM TTR (Control 2). A fixed volume (≈ 20 – 30 nL) of Control 1 (Trace A & B) and Control 2 (Trace C & D) was injected and separated by CZE in duplicates. The reduction in free 8-ANS peak height/area indicates 8-ANS binding to TTR in Control 2.

Based on the peak heights or areas of free 8-ANS with/without TTR, it can be deduced that around 60% or 35% of 8-ANS was bound to TTR at the specified concentration ratio. Significant discrepancy in the deduced fractional occupancy of TTR was due to peak tailing and broadening effect caused by molecular binding which inflates peak area but not height value demonstrated later when compounds were introduced into the Running Buffer. The percentage of 8-ANS bound TTR calculated from peak height also agrees better with previously reported K_d_ values (1 – 5 µM) [28], henceforth, free 8-ANS peak height was the chosen measurement parameter. With 20 μM of TTR and two binding sites per tetramer, a minimum of 40 μM of 8-ANS is required to occupy all sites. A 60% reduction in free 8-ANS peak height indicates that around 18 μM of 8-ANS was bound to TTR, *i.e.* most of the TTR were singly bound by 8-ANS. The free 8-ANS (30 μM) peak height from Inject Buffer without TTR is now defined as Control 1 Peak Height, and Control 2 Peak Height is from the 8-ANS (30 μM)/TTR (20 μM) mixture. From subsequent multiple runs of Control 1 & 2 throughout assay development, peak height reproducibility and its derived Z′-factor was far better (**Table 1**) than that of area, further consolidating the use of peak height.

**Table 1.** Reproducibility of Free 8-ANS Peak Height/Area Values of Control 1 and 2

| **Number of Experiments = 16** | **Peak Height** | **Peak Area** |
| --- | --- | --- |
| Control 1 Average (N = 46) <sup>a</sup> | 1586.57<br>95% CI [1548.15 – 1624.99] | 12421.03<br>95% CI [11911.86 – 12930.20] |
| Control 2 Average (N = 102) <sup>a</sup> | 505.01<br>95% CI [499.00 – 511.02] | 5867.50<br>95% CI [5699.86 – 6035.14] |
| Control 1 Intra-Assay Deviation <sup>b</sup> | Lowest 2.12 (0.15%)<br>Highest 123.04 (8.47%)<br>Average 2.72% | Lowest 130.11 (1.16%)<br>Highest 1772.05 (14.18%)<br>Average 6.46% |
| Control 2 Intra-Assay Deviation <sup>b</sup> | Lowest 4.58 (0.97%)<br>Highest 44.86 (8.97%)<br>Average 2.85% | Lowest 185.17 (3.74%)<br>Highest 967.90 (16.88%)<br>Average 7.29% |
| Control 1 Inter-Assay Deviation <sup>c</sup> | 132.94 (8.38%) | 1761.95 (14.19%) |
| Control 2 Inter-Assay Deviation <sup>c</sup> | 30.95 (6.13%) | 863.85 (14.72%) |
| Z'-factor <sup>d</sup> | 0.54 | -0.20 |

<sup>a</sup> The average of free 8-ANS peak height/area values measured from 4 – 5 different Inject Buffer vials prepared on 4 – 5 different dates and separated in 3 different coated capillaries, see **S1 File** for details.
<sup>b</sup> The range of standard deviation (SD) and coefficient variation in parenthesis of measured free 8-ANS peak height/area. The repetitive CZE runs were performed from the same vial, on the same date, with the same coated capillary and buffer batch.
<sup>c</sup> The standard deviation and coefficient variation in parenthesis of measured free 8-ANS peak height/area from a total of 46 (Control 1) and 102 (Control 2) replicate runs.
<sup>d</sup> Z' factor [40] was calculated by treating free 8-ANS peak height/area of Control 1 as the positive and Control 2 as the negative.

### Reproducibility of Control 1 & Control 2 Peak Heights

To become a sensitive assay, the free 8-ANS peak height or area of Control 1 and 2 must be consistent so that any increment could be accounted to 8-ANS displacement from TTR. A fixed volume of Control 1 and Control 2 was injected and separated numerous times in different occasions. **Table 1** summarises the inter-assay and intra-assay deviation of Control 1 & 2 free 8-ANS peak height values. They were highly reproducible with a Z′-factor of 0.54, supporting the basis for a reliable fragment screening assay. Based on values from **Table 1**, the cut-off for a positive hit was set to a minimum of 10% free 8-ANS peak height restoration equivalent to > 3σ of the inter-assay Control 2 deviation value.

### Effect of Competitors in the Running Buffer on Free 8-ANS Peak Height

The next step in a typical ACE assay would be to add competitors into the Inject Buffer containing 8-ANS and TTR and observe the change in free 8-ANS peak height or area. This however will significantly increase protein consumption because the mixture cannot be reused to test the next compound, especially if [protein] were at micromolar level with sample volumes set to tens of microlitres. Fluorescence detection could be employed to lower protein concentration used for screening, but it is not always possible/ideal due to various reasons. To avoid mixing the 8-ANS/TTR solution with test ligand, the author devised an alternative method (the FPPHR method) in which the competitor compound was added to the Running Buffer instead of Inject Buffer. The injected 8-ANS-TTR mixture were separated in a Running Buffer containing the test compound. Therefore, the same vial of Inject Buffer could be used throughout the screen and only 20 – 30 nL of the protein solution is consumed for each run. This means 100 µL of the 30 µM 8-ANS 20 µM TTR mixture solution is enough for around 2500 – 3000 screens, using just 0.11 mg of TTR protein.

To see if 8-ANS could be displaced from the thyroxine binding sites of TTR, natural ligand Thyroxine T_4_, Tafamidis, and a few substructures of a known TTR ligand [24] were selected for testing. Thyroxine T_4_ (**Fig 3A**), 3,5-Dichloroaniline (**Fig 3B**, **Trace C – H**), 4-Amino-2,6-Dichlorophenol (**S3 Fig A, Trace B),** Diphenylamine (**S3 Fig B, Trace A**), and Tafamidis (**S4 Fig A, Trace B)** all led to significant free 8-ANS peak height increase. To ensure assay fidelity, compounds expected not to bind to TTR were also tested. The free 8-ANS peak height changed minimally when 2-Aminobenzenesulphonamide (**S3 Fig A, Trace A**) Verapamil (**S3 Fig B, Trace B**), or Acetazolamide (**S4 Fig A, Trace A**) were added to the Running Buffer.

**Fig 3.**
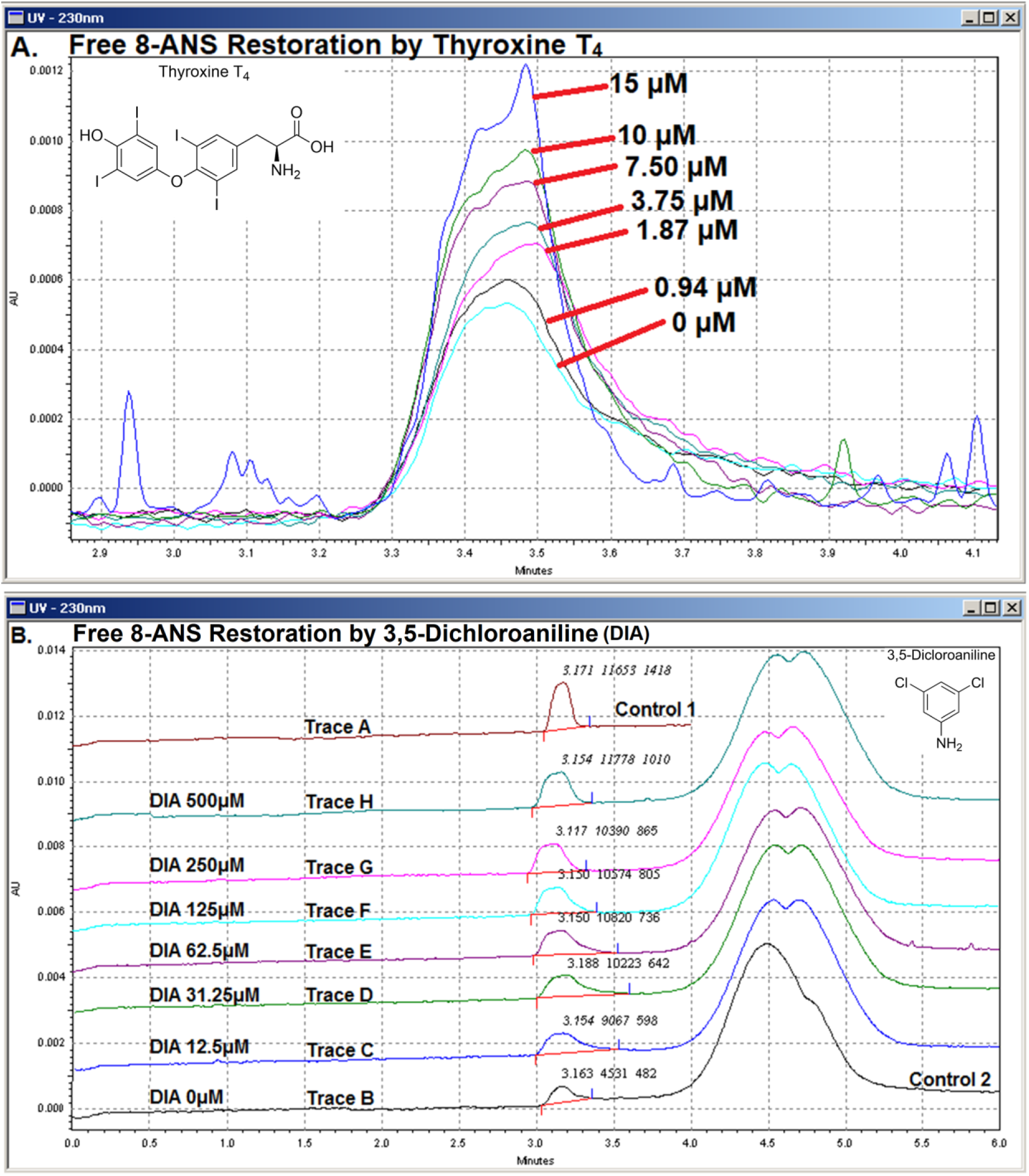
CZE Assay Based on Free 8-ANS Peak Height Restoration. (**A.**) A sequence of CZE separation runs was performed with increasing concentration of Thyroxine T4 added to the Running Buffer (20 mM HEPES pH 7.0, 2 mM CaCl2, 1% v/v DMSO f/c). A fixed volume of Inject Buffer (containing 30 μM 8-ANS and 20 μM TTR) was injected into the capillary filled with Running Buffer. The free 8-ANS UV peaks were superposed and zoomed in, omitting the absorbance peak of TTR that elutes later. Evidently, free 8-ANS peak height increased in response to increasing concentration of Thyroxine T4 in the Running Buffer. The baseline was particularly noisy when [Thyroxine T4] reaches 15 µM, which might be caused by its limited aqueous solubility. (**B.**) A fixed volume of Control 1 (Trace A) & Control 2 (Trace B) was injected and separated. Control 2 injection and separation was repeated but with increasing concentration of 3,5-Dichloroaniline (DIA) added to the Running Buffer (Trace C – H). Free 8-ANS peak height and area increased accordingly meaning DIA was displacing 8-ANS from TTR. Migration time of free 8-ANS and TTR protein were different to that of **Fig 2** because a different coated capillary was used. Numeric values atop free 8-ANS peaks indicate migration time (minutes), peak area and peak height in arbitrary units (AU) respectively.

It is worth noting that restored free 8-ANS peak area changed very little with increasing concentration of 3,5-Dichloroaniline after 12.5 µM yet peak height restoration continued steadily (**Fig 3B**, **Trace C – H**). As for the negative control compounds, free 8-ANS peak area jumped by 14 – 24 % with minimal peak height increase (**S3 and S4 Fig**). These observations suggest that peak area is more susceptible to peak shape distortion due to the complex molecular binding competition events that took place during native electrophoresis. The early saturation of peak area restoration by 3,5-Dichloroaniline presents another reason for it not being used to monitor 8-ANS displacement. In any event, sensitive binding detection of fragment compound 3,5-Dichloroaniline by the short-contact-time FPPHR method has emboldened its utilisation in fragment screening.

### TTR CZE FPPHR Fragment Screening and Hit Validation by Crystallography

Having ascertained reproducibility of control values (sensitivity) and reliability in binder identification (specificity), 129 fragments from the Selcia Fragment Library (SFL) were screened at 300 µM using the FPPHR method. 16 fragments (12.4 % hit rate) have restored ≥ 10% of Free 8-ANS Peak Height value (**Table 2**). Titration experiments were carried out on 15 (fresh DMSO stocks dissolved from solids) of the 16 initial fragment hits, they all led to a dose-dependent increment of free 8-ANS peak height (**S2 File**) like seen in **Fig 3**. Dose-response of one of the weakest detectable fragments SFL000006 – 3,5-Dichlorobenzenesulfonamide (DSF) is shown in **S4 Fig B**.

**Table 2.**
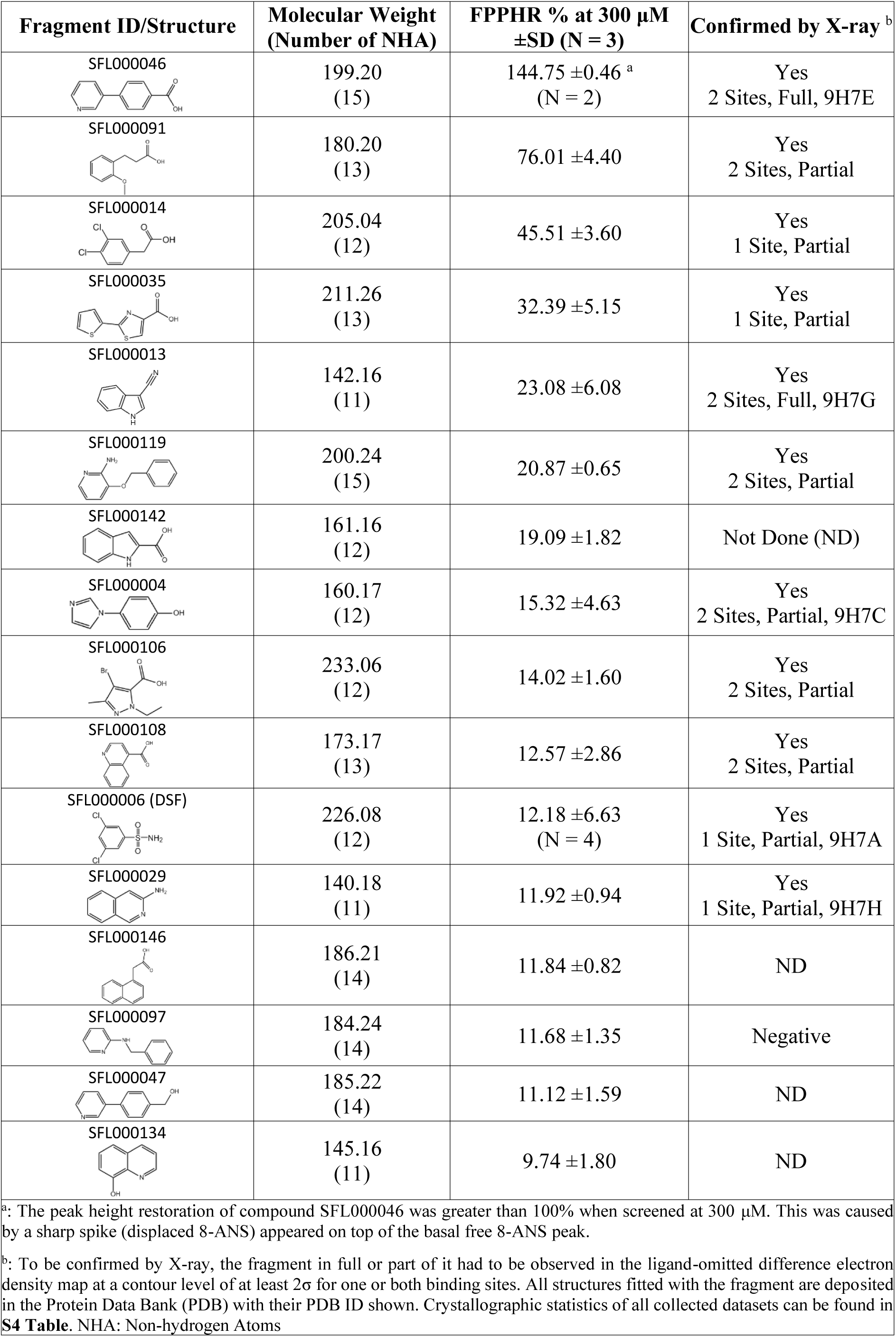
TTR CZE FPPHR Fragment Hits

| Fragment ID/Structure | Molecular Weight<br>(Number of NHA) | FPPHR % at 300 μM<br>±SD (N = 3) | Confirmed by X-ray<sup>b</sup> |
| --- | --- | --- | --- |
| SFL000046 | 199.20<br>(15) | 144.75 ±0.46<sup>a</sup><br>(N = 2) | Yes<br>2 Sites, Full, 9H7E |
| SFL000091 | 180.20<br>(13) | 76.01 ±4.40 | Yes<br>2 Sites, Partial |
| SFL000014 | 205.04<br>(12) | 45.51 ±3.60 | Yes<br>1 Site, Partial |
| SFL000035 | 211.26<br>(13) | 32.39 ±5.15 | Yes<br>1 Site, Partial |
| SFL000013 | 142.16<br>(11) | 23.08 ±6.08 | Yes<br>2 Sites, Full, 9H7G |
| SFL000119 | 200.24<br>(15) | 20.87 ±0.65 | Yes<br>2 Sites, Partial |
| SFL000142 | 161.16<br>(12) | 19.09 ±1.82 | Not Done (ND) |
| SFL000004 | 160.17<br>(12) | 15.32 ±4.63 | Yes<br>2 Sites, Partial, 9H7C |
| SFL000106 | 233.06<br>(12) | 14.02 ±1.60 | Yes<br>2 Sites, Partial |
| SFL000108 | 173.17<br>(13) | 12.57 ±2.86 | Yes<br>2 Sites, Partial |
| SFL000006 (DSF) | 226.08<br>(12) | 12.18 ±6.63<br>(N = 4) | Yes<br>1 Site, Partial, 9H7A |
| SFL000029 | 140.18<br>(11) | 11.92 ±0.94 | Yes<br>1 Site, Partial, 9H7H |
| SFL000146 | 186.21<br>(14) | 11.84 ±0.82 | ND |
| SFL000097 | 184.24<br>(14) | 11.68 ±1.35 | Negative |
| SFL000047 | 185.22<br>(14) | 11.12 ±1.59 | ND |
| SFL000134 | 145.16<br>(11) | 9.74 ±1.80 | ND |

<sup>a</sup>: The peak height restoration of compound SFL000046 was greater than 100% when screened at 300 μM. This was caused by a sharp spike (displaced 8-ANS) appeared on top of the basal free 8-ANS peak.
<sup>b</sup>: To be confirmed by X-ray, the fragment in full or part of it had to be observed in the ligand-omitted difference electron density map at a contour level of at least 2 $\sigma$ for one or both binding sites. All structures fitted with the fragment are deposited in the Protein Data Bank (PDB) with their PDB ID shown. Crystallographic statistics of all collected datasets can be found in **S4 Table**. NHA: Non-hydrogen Atoms

12 of the 15 titrated hits alongside 3,5-Dichloroaniline (DIA) and N-Phenylanthranilic Acid (NPA) were selected for co-crystallisation with TTR (**S5 Fig A – C**). 11 of the 12 fragments were confirmed (92 % confirm rate) by either partial or full ligand electron density occupation in one or both binding sites (**Table 2**). Other than hit-validation, crystal structure of DIA and DSF in complex with TTR has highlighted the precariousness in the binding mode of fragment analogues. The increased degree of freedom for fragments in the binding site has made prediction more difficult. However, when fragments are optimised into part of a lead compound, they do not seem to change the way they bind to the target. This is illustrated by the superposition of DIA and NPA structure to that of a previously characterised TTR ligand 2-(3,5-dichloro-4-hydroxyphenylamino)benzoic acid (DHB) [24], their counterpart atomic positions accurately matched (**S5 Fig D**).

Two chemical series were identified from the 16 initial hits, carboxylate substituted aromatic rings and fused heterocycles, of which the former is known to bind TTR promiscuously [41]. In pursuit of chemical novelty, an analogue screen using immediately available SFL compounds was conducted (**S1 Table**) on fragment hit SFL000013 leading to discovery of the 4-(3H-pyrazol-4-yl)quinoline (SFL001535) scaffold.

### Follow-up Studies of SFL000013 & SFL001535

#### Structural Characterisation of SFL000013 and SFL001535

Being a grown version of SFL000013, and with marked increase in FPPHR %, the two compounds were co-crystallised with TTR. SFL00013 occupied the inner part of the binding channel, of which the indole nitrogen formed a hydrogen bond with the Ser117 sidechain (**Fig 4A**). Intriguingly, SFL001535 binds to the TTR tetramer in two modes (**Fig 4B**). Its pyrazole nitrogen hydrogen bonds with Ser117 whilst the quinoline sits in the mid part of the binding channel in Mode A and flipped 180° horizontally having the quinoline nitrogen forms hydrogen bond with Ser117 in Mode B. This dual binding mode of SFL001535 remained for the aggressively amyloidogenic S52P [42] TTR mutant in cocrystal structure. Although binding kinetics and dynamics are usually not observable in crystal structures, the SFL001535 binding modes here have structurally demonstrated binding cooperativity.

**Fig 4.**
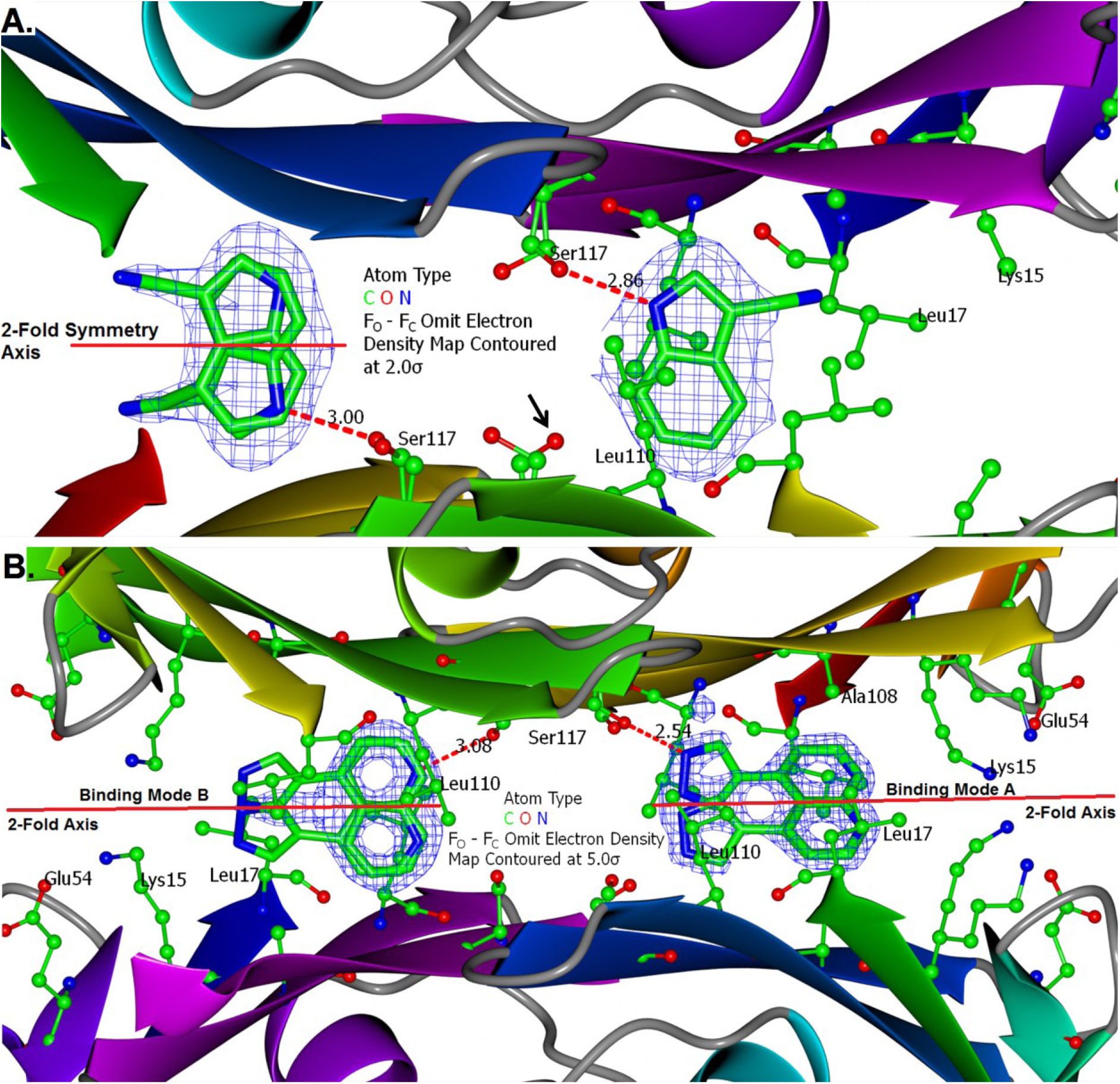
Binding of SFL000013 and SFL001535 to TTR. (**A.**) Binding of SFL000013 to TTR involved a hydrogen bond between the 1*H*-Indole and the hydroxyl of Ser117. The fused aromatic ring fits in between two Leucine sidechains (Leu110) from two TTR monomers. Electron density for the cyano group is less defined, but it has strengthened the 1*H*-Indole hydrogen bond with Ser117 as suggested by FPPHR assay data in **Table S1**. The unengaged Serine (black arrow) is potentially available for additional hydrogen bonds. (**B.**) SFL001535 was bound to TTR in two distinct modes. The pyrazole formed a strong (bond length = 2.54 Å) hydrogen bond with Ser117 in Mode A. Extensive hydrophobic contacts were made between the quinoline and the two Leucine sidechains (Leu17) from two TTR monomers. In Mode B, SFL001535 has flipped around and the quinoline is positioned where the indole of SFL000013 was. Although the two binding sites looked identical in crystal structures, binding of SFL001535 has provided ligand-based structural evidence for cooperative binding in TTR.

#### Induced Circular Dichroism (CD) of SFL001535

In addition to ACE and crystallography, binding of SFL001535 to TTR was further investigated by induced circular dichroism. When an achiral molecule SFL001535 (**Fig 5A**, **Red Circles**) binds to a chiral molecule TTR (**Fig 5A**, **Blue Squares**), the UV absorption (280 – 320 nm) of SFL001535 will become circularly dichroic. This is revealed (**Fig 5B**) when the CD spectrum of TTR mixed with SFL001535 is subtracted by CD spectrum of TTR alone. Though extra insight into the mechanism of cooperative binding was not gained, their binding is nevertheless confirmed again by a second orthogonal assay.

**Fig 5.**
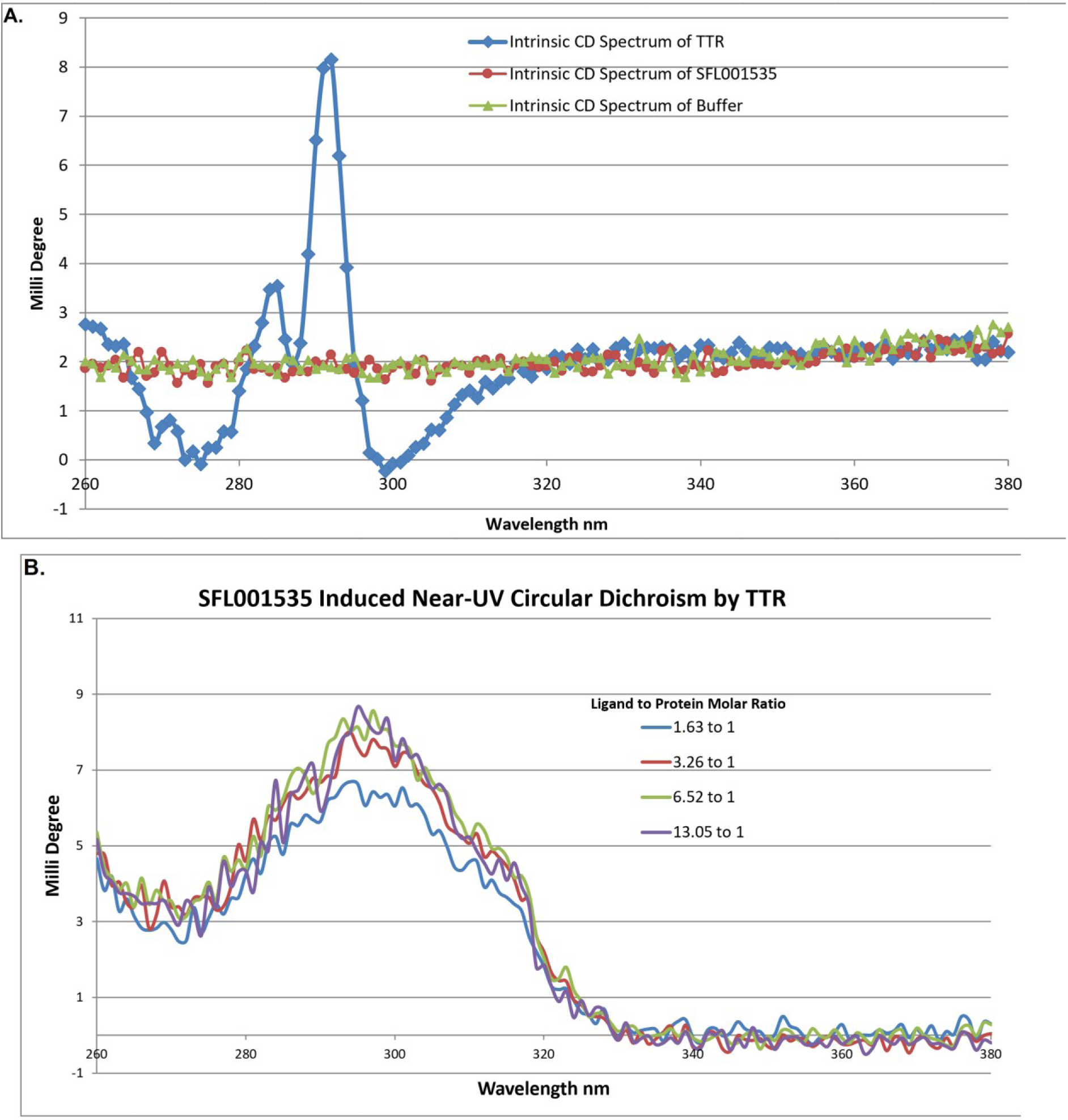
Near-UV CD Analysis of SFL001535 Binding to TTR. (**A.**) There was no intrinsic near-UV CD spectrum observed for the buffer solution and compound SFL001535. The intrinsic near-UV CD spectrum of TTR is shown as two peaks at around 285 nm and 292 nm with a trough before and after those peaks. (**B.**) A near-UV CD spectrum scan was performed on a mixture containing TTR and compound SFL001535 in buffer solution. Having subtracted the intrinsic CD signal of TTR, a peak spanning from 280 – 320 nm was observed. The peak represents the induced CD signal of SFL001535 upon TTR binding. The signal intensity at 300nm became stronger as the ligand-to-protein molar ratio increased from 1.62 (blue) to 3.26 (red). However, there was little change with further increase of ligand concentration (green & purple). It was likely that all TTR binding sites were occupied by the ligand at an excess of 3.26 to 1.

#### Structure-Activity Relationship (SAR) Build-up

Guided by gathered structural data on SFL000013 and SFL001535, more readily available compounds were selected for testing by FPPHR from sources such as the Selcia Fragment Library, Selcia Chemical Store, and commercial vendors to build up SAR. The crystal structures (**Fig 4**) including that of SFL001561 (**S6 Fig**) and results from **S1 and S2 Tables** have yielded a detailed perspective on the binding of aromatic heterocycles to TTR. Firstly, the inner part of the binding channel near Ser117 averts from increased basicity and reduced hydrophobicity. Secondly, hydrogen bonding between Ser117 sidechain and heterocyclic nitrogen atoms is enhanced by electron-withdrawing groups substitution and weakened by electron donors. Preference of Ser117 sidechain to act as hydrogen bond acceptor was further emphasised with breakage of pyrazole tautomer in compound SFL001562 and its severe loss of FPPHR activity (**S2 Table**). Finally, the mid-part of the binding channel formed by the Leu17 and Ala108 sidechains felicitously accommodates hydrophobic aromatic rings.

Based on the SAR data and crystal structures obtained, five lead molecule scaffolds were proposed (**Fig 6**). Scaffold A is to grow SFL001535 as crystal structure has revealed there are unfilled spaces available on position 3 and 5 of the pyrazole ring. SAR data from CZE assays suggests that R_1_ and R_2_ substituents should be electron withdrawing and/or hydrophobic. Effect of the location of quinoline nitrogen on compound binding is however still unknown. The second stage of ligand growth could be to add R_3_ substituents onto the quinoline at position 16, which could form additional ionic interactions with Lys15 and Glu54 sidechain, see **Fig 4B**. Scaffold B – E are the merged variants of SFL00013 and SFL001535. The double fused-ring system is constructed by the attachment of quinoline onto the diazole part (Scaffold B) or the benzene part of indazole (Scaffold C). Both scaffolds are predicted to bind to TTR with the indazole near to Ser117 and quinoline midway of the binding channel. R_1_ substituents are designated to be electron-withdrawing groups that would enhance hydrogen bond between the 1H-Indazole and Ser117 sidechain. R_2_ substituents should be hydrogen bond donors for the other Ser117 sidechain on another TTR monomer. Scaffold D is drawn to circumvent some probable steric hindrance problems associated with Scaffold B & C. By replacing the quinoline with a phenyl, there would be a lot more room for the R_1_ substituent though preference for R_3_ substituents remains unclear. Finally, the dihydropyrazolopyrazole (Scaffold E) is expected to form hydrogen bonds to two Ser117 sidechains from two TTR monomers, hence R_2_ substituent is not needed. Its smaller size compared to indazole should also provide a minor relief on rotational constraint of the quinoline.

**Fig 6.**
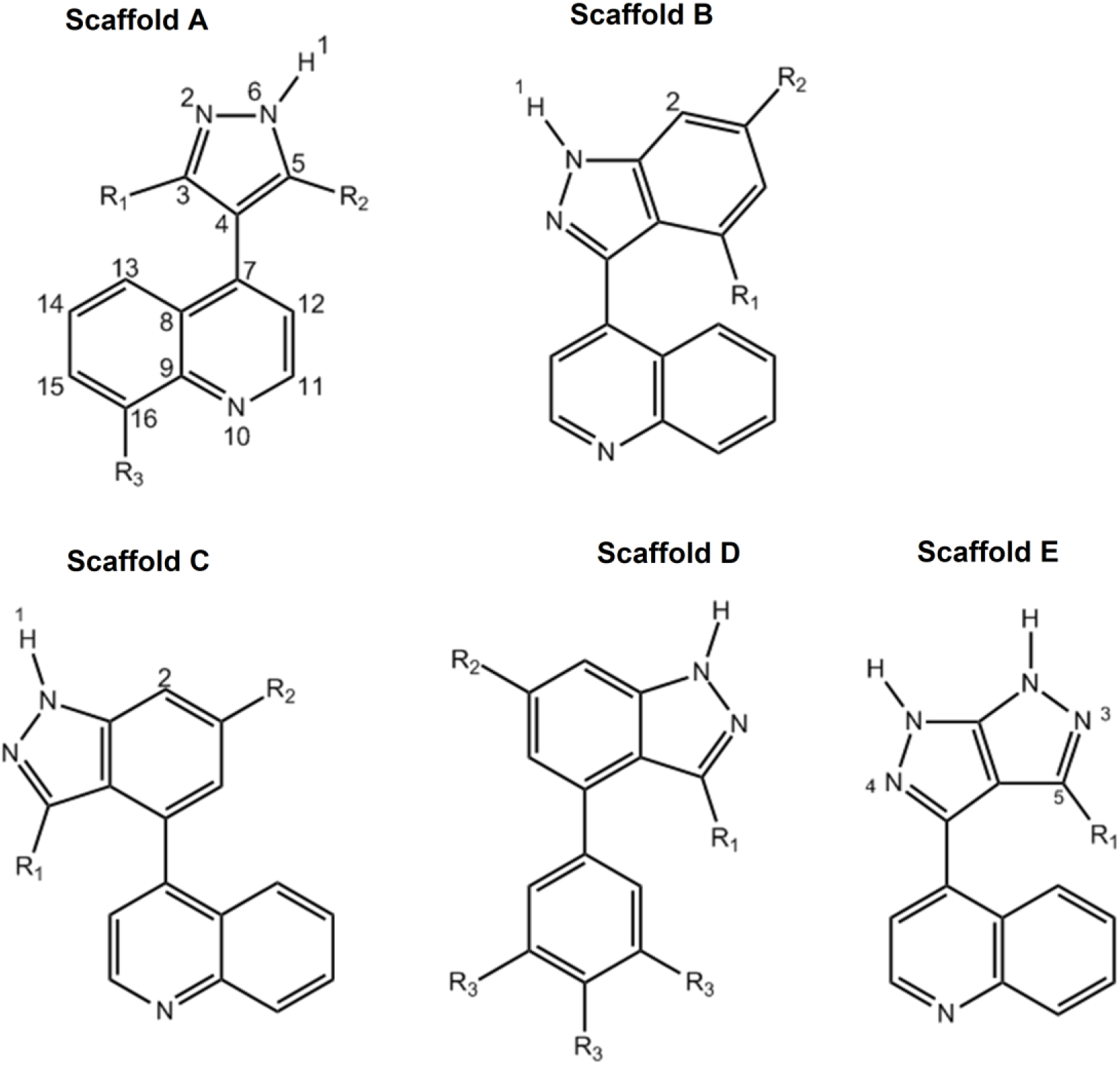
Proposed Heterocycle-based TTR Kinetic Stabilisers

Close analogues of the proposed leads (Scaffold B – D) were purchased from commercial vendors and tested by FPPHR (**S3 Table**) and some co-crystallised with TTR. Scaffold A remained superior to the others and focus was shifted to growing SFL001535, leading to the discovery of Compound 94320297 and synthesis of SEL101928. Unlike SFL001535, Compound 94320297 binds to TTR in one mode only and the quintessential direct linkage between pyrazole and quinoline has allowed the ligand to sterically optimise its binding with Ser117, Leu110, Leu17 and Ala108 (**Fig 7**).

**Fig 7.**
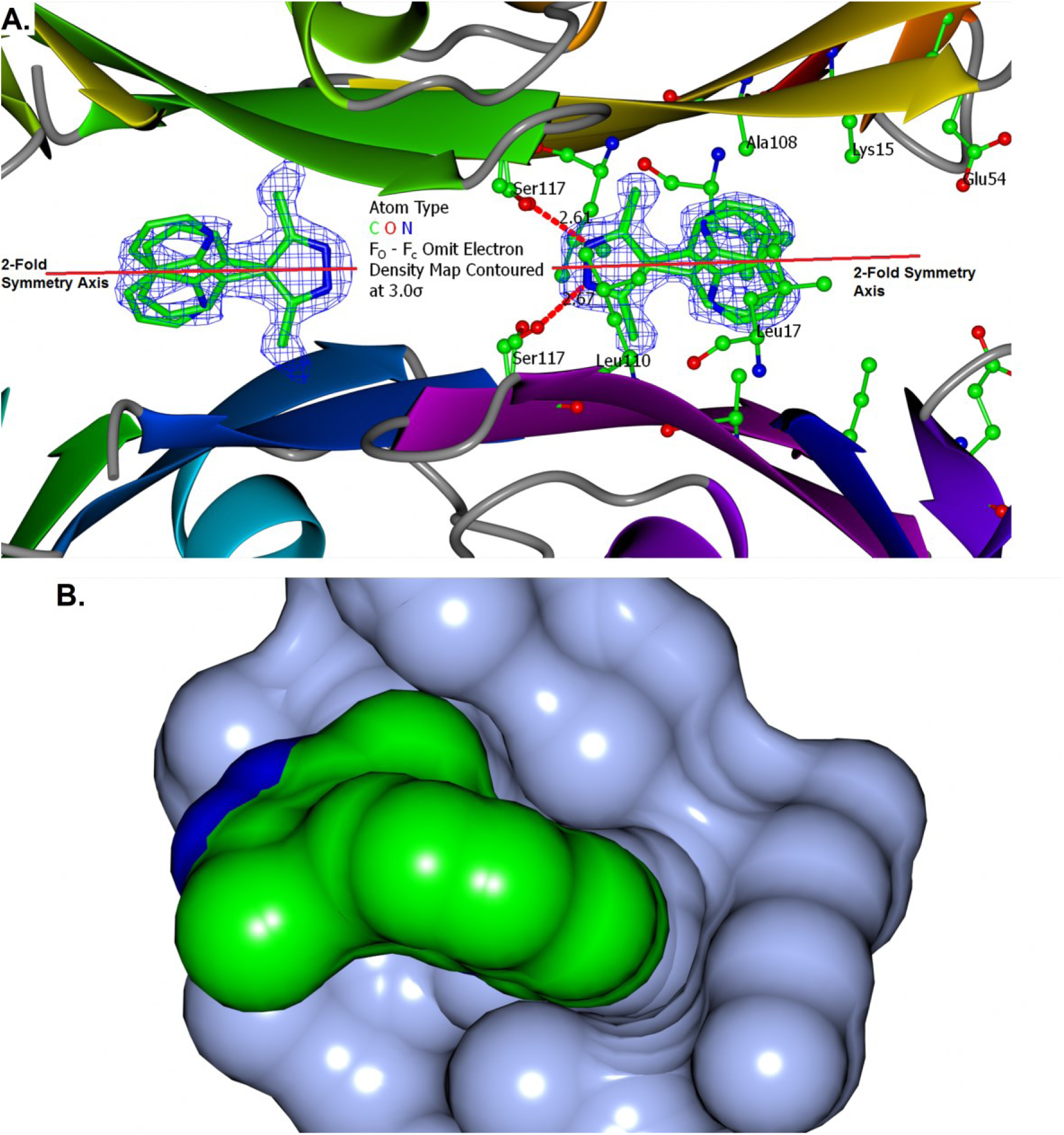
TTR and Compound 94320297 Binding. (**A.**) In contrast to SFL001535, Compound 94320297 binds to TTR with a single binding mode. The dimethylpyrazole positioned its centre along the 2-fold axis and hydrogen bonded to two Ser117 sidechains. (**B.**) As well as the special double-Ser117 hydrogen bond configuration, the quinoline of 94320297 is also a major affinity contributor. A surface display shows it sitting tilted in a prime location for extensive hydrophobic/vdW contacts sandwiched in between two Leu17 sidechains.

#### Isothermal Titration Calorimetry (ITC)

To determine accurately the TTR binding affinity of lead molecules derived from Scaffold A, Compound SFL001535, 94320297, SEL101928 and Tafamidis were subjected to ITC measurement. Binding isotherm of Tafamidis appeared to be sigmoidal (**Fig 8A**) and one-site model (no binding cooperativity) provided a better fit of the data resulting a single K_d_ of 229 nM. This is discrepant to its previously reported [43] ITC K_d_ of 3 nM and 278 nM. Negative cooperativity was evident from the ITC isotherms of the other tested compounds (**Fig 8 B – D**). Nonetheless, Compound SEL101928 and 94320297 had comparable K_d_ values to that of Tafamidis (**Table 3**).

**Fig 8.**
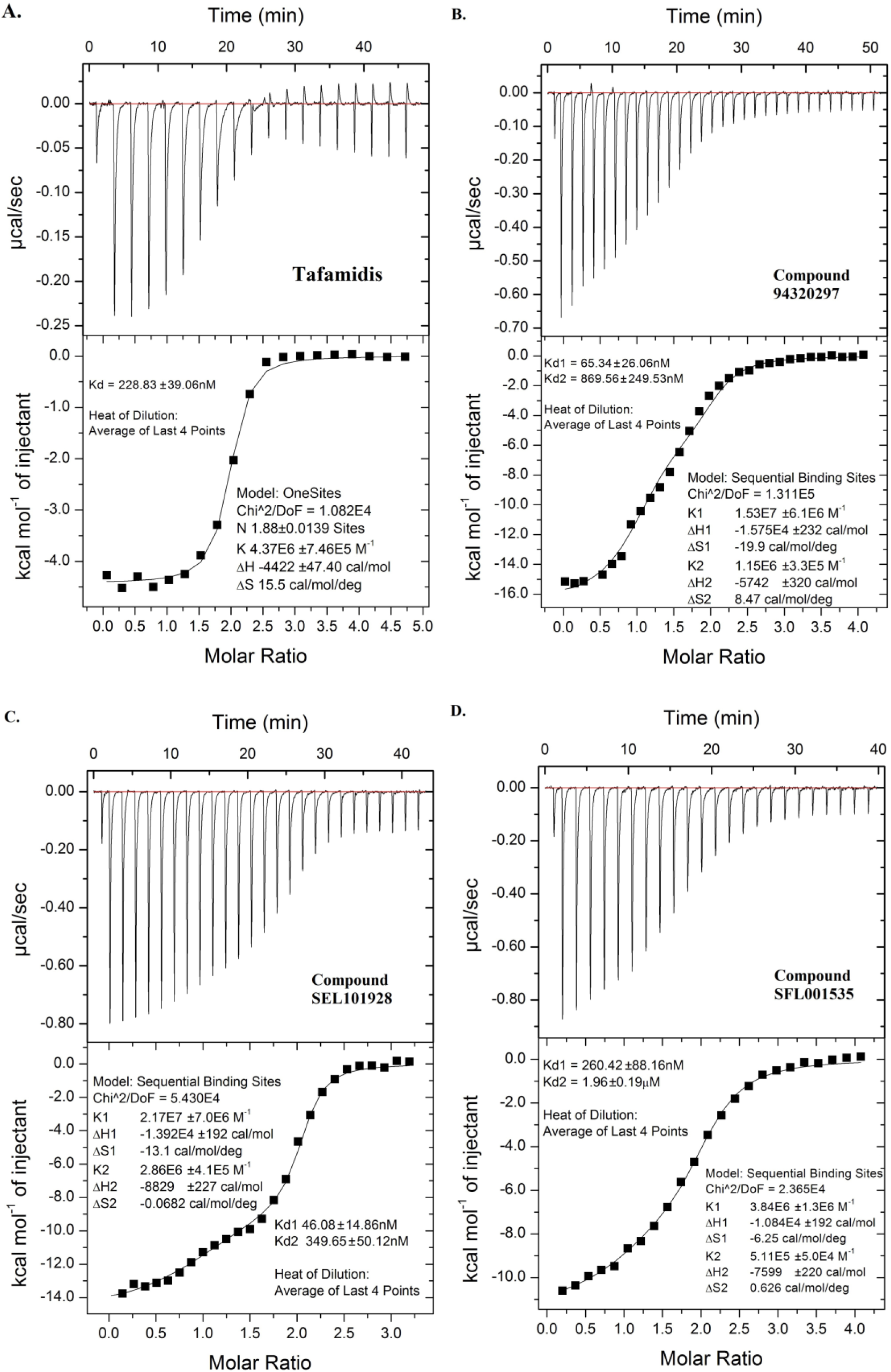
ITC Binding Isotherms of Tafamidis, SFL001535, 94320297 and SEL101928. Isotherms of tested compounds along with calculated Kd values and standard error of curve fitting are shown. Isotherm of Tafamidis appeared to be sigmoidal (**A.**) whilst binding cooperativity was evident for Compound 94320297 (**B.**), SEL101928 (**C.**) and SFL001535 (**D.**)

**Table 3.**
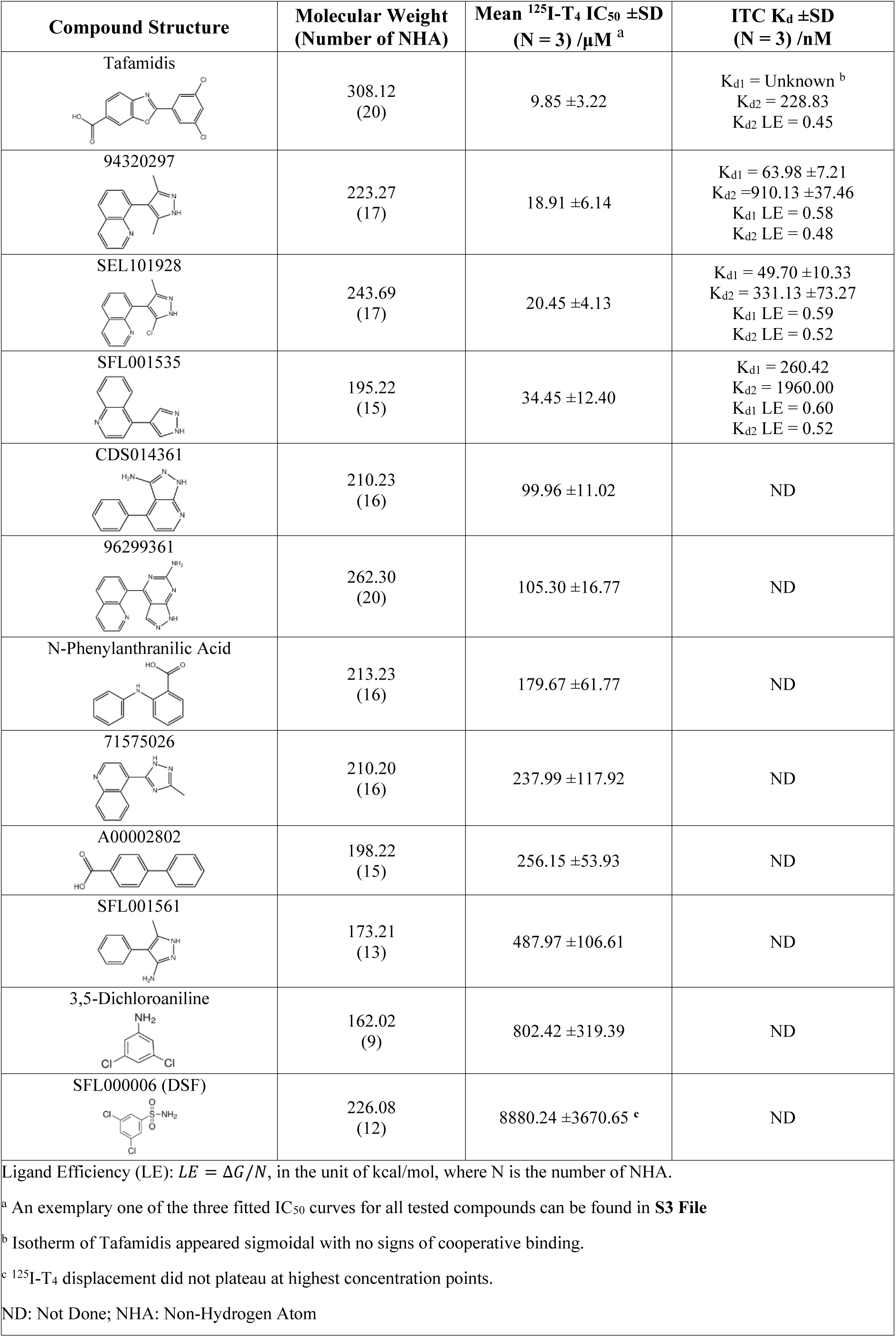
Isothermal Titration Calorimetry and ^125^I-T4 Displacement Plasma Binding Assay

| Compound Structure | Molecular Weight<br>(Number of NHA) | Mean <sup>125</sup>I-T<sub>4</sub> IC<sub>50</sub> ±SD<br>(N = 3) /μM <sup>a</sup> | ITC K<sub>d</sub> ±SD<br>(N = 3) /nM |
| --- | --- | --- | --- |
| Tafamidis | 308.12<br>(20) | 9.85 ±3.22 | K<sub>d1</sub> = Unknown <sup>b</sup><br>K<sub>d2</sub> = 228.83<br>K<sub>d2</sub> LE = 0.45 |
| 94320297 | 223.27<br>(17) | 18.91 ±6.14 | K<sub>d1</sub> = 63.98 ±7.21<br>K<sub>d2</sub> = 910.13 ±37.46<br>K<sub>d1</sub> LE = 0.58<br>K<sub>d2</sub> LE = 0.48 |
| SEL101928 | 243.69<br>(17) | 20.45 ±4.13 | K<sub>d1</sub> = 49.70 ±10.33<br>K<sub>d2</sub> = 331.13 ±73.27<br>K<sub>d1</sub> LE = 0.59<br>K<sub>d2</sub> LE = 0.52 |
| SFL001535 | 195.22<br>(15) | 34.45 ±12.40 | K<sub>d1</sub> = 260.42<br>K<sub>d2</sub> = 1960.00<br>K<sub>d1</sub> LE = 0.60<br>K<sub>d2</sub> LE = 0.52 |
| CDS014361 | 210.23<br>(16) | 99.96 ±11.02 | ND |
| 96299361 | 262.30<br>(20) | 105.30 ±16.77 | ND |
| N-Phenylanthranilic Acid | 213.23<br>(16) | 179.67 ±61.77 | ND |
| 71575026 | 210.20<br>(16) | 237.99 ±117.92 | ND |
| A00002802 | 198.22<br>(15) | 256.15 ±53.93 | ND |
| SFL001561 | 173.21<br>(13) | 487.97 ±106.61 | ND |
| 3,5-Dichloroaniline | 162.02<br>(9) | 802.42 ±319.39 | ND |
| SFL000006 (DSF) | 226.08<br>(12) | 8880.24 ±3670.65 <sup>c</sup> | ND |

Ligand Efficiency (LE): $LE = \Delta G/N$ , in the unit of kcal/mol, where N is the number of NHA.
<sup>a</sup> An exemplary one of the three fitted IC<sub>50</sub> curves for all tested compounds can be found in **S3 File**
<sup>b</sup> Isotherm of Tafamidis appeared sigmoidal with no signs of cooperative binding.
<sup>c</sup> <sup>125</sup>I-T<sub>4</sub> displacement did not plateau at highest concentration points.
ND: Not Done; NHA: Non-Hydrogen Atom

## ^125^I-T_4_ Displacement in Neat Normal Plasma

Besides binding to TTR in aqueous buffer *in vitro*, potential biological function of the lead molecules must be verified to improve the likelihood of future clinical efficacy. A first step is to ensure binding of the lead molecules to TTR remains significant in the presence of all other blood proteins, albumin in particular. This was tested by displacement of radiolabelled thyroxine ^125^I-T_4_ from TTR in neat normal plasma. A selection of compounds was chosen, and all were found to displace Thyroxine T_4_ in the assay with different potency (**Table 3**). In good correlation to ITC K_d_ values, lead compound SEL101928 and 94320297 had similar IC_50_ values to that of Tafamidis, highlighting their potentials in becoming an effective TTR kinetic stabiliser.

## Discussion and Conclusions

In a CE separation apparatus, the user has full control on the contents of Running Buffer, Inject Buffer and filling of the capillary. This assay setup flexibility has facilitated development of different modes of separation and numerous variants of ACE methods including the patented CEfrag™ method. FPPHR materially differs from them by not filling the capillary with protein-containing solution and not mixing test compounds with the protein. Not only this will reduce protein consumption but also lessen the issue of protein adsorption onto the capillary surface, making a single capillary to last the entire screen possible. Protein consumption is further reduced by its non-mixing format and reusing the protein-probe ligand solution (Inject Buffer). In our case, 3000 fragments can be screened with just 0.11 mg of TTR protein.

Further to low protein consumption, the FPPHR method simplifies assay procedure. Multiple Running Buffers containing the screened fragment at a fixed concentration could be prepared in just two pipetting steps, which is the addition of fragment DMSO stock and its dilution with Buffer B. Although an automatable procedure, limited by the design of the instrument, only ≈ 30 pairs of Running Buffers could be loaded on the buffer tray and changing the next set would require manual input. Considering a typical total run time of 6 – 8 minutes per fragment, around 60 – 90 fragments could be screened per worker per machine on a 9-5pm shift. This throughput might be unsatisfactory for some, especially in the industry sector. However, contrary to HTS, throughput becomes less of a priority in FBDD, partly because the fragment library size (0.5 – 2k) is much smaller, but also the pivot is on hit quality and reliability. Potentially, throughput could be increased through the instrument manufacturer by increasing the buffer tray capacity and installing a capillary array.

Apart from concerns on screening throughput, the FPPHR method might be less discerning on high affinity binders. For example, the 8-ANS displacement was expected to be more significant with 10 µM of Tafamidis (**S4 Fig A, Trace B**) and it was unable to reflect the binding affinity difference between SFL001535 and 94320297 (**S2 and S3 Tables**). A probable explanation for the occasional loss of assay fidelity is the circumstances in which 8-ANS displacement occurs. The competitor would only meet up with 8-ANS bound TTR after on-column injection *i.e.* the non-mixing nature of FPPHR method. The injection plug (≈ 30 nL) is then sandwiched between the Running Buffer containing the competitor in front inside the capillary and behind from the outlet reservoir (**Fig 1**). Electrophoresis starts soon (≈ 10 s) after the injection step, leaving little incubation time between the competitor, 8-ANS and TTR. Their competitive binding might not have reached equilibrium prior to electrophoresis, especially for binders with exceptionally slow K_on_. Even so, FPPHR excels in picking up binding fragments in a yes/no manner, which fits the purpose of a primary screening tool. Standard binding assays such as ITC should then be used to accurately determine binding affinity at equilibrium of fragment-derived lead molecules. Moreover, short exposure time for competitive displacement intuitively implies Probe Ligands of lower affinity is preferred.

Subsequently, molecular interactions that occur during an electrophoretic event in a non-denaturing environment could be tremendously intricate. Once the voltage is on, all species including neutral molecules inside the capillary are moving under the influence of electromotive force. We know 8-ANS and TTR were migrating towards the positive terminal because they are negatively charged. However, the electrophoretic velocity of test compounds is variable depending on their structure and net charge. The way how the competing compound encounters the 8-ANS-TTR complex during electrophoresis could be another contribution factor to 8-ANS displacement besides binding affinity. The detailed molecular events that occurred at the interface between the injection plug and the Running Buffer immediately prior and during electrophoresis would require future elucidation.

The developed FPPHR method here is a reproducible, microscale, non-mixing, and solution-based CZE separation process using physiological buffer with sensitive UV detection in a competition format. As well as sensitive binding detection and low protein consumption, those characteristics conferred additional distinctive advantages to its utilisation in fragment screening. Firstly, it is site-specific searching for fragment that bind to the site of interest only. Secondly, the binding events occur in a free solution environment where the natural movement of the drug target is not restrained nor immobilised. This is particularly important for dynamic targets that exhibit conformational change or subunit movement upon ligand binding. Thirdly, CZE is particularly responsive to compound precipitation events in aqueous buffer, which would fail the electropherogram but could result in false positives in other screening techniques. Having this built-in filter for insoluble, intractable compounds could help save time from making futile attempts. Lastly, its UV detection means the Probe Ligand and drug target can be as native as possible without any labelling. Together with the purifying effect of CZE, FPPHR will be less prone to fluorescence and high concentration interference.

From the process of CZE fragment hit validation by crystallography, a few interesting observations were also made. Although molar excess of DIA and DSF to TTR were 235x in the crystallisation drop, yet their electron density was only observed in one site, providing ligand-based structural evidence for negative cooperative binding. Another observation is the precise atomic position matching of DIA and NPA to their merged form DHB (**S5 Fig D**). The structures of DIA and NPA in complex with TTR have correctly predicted the atomic position of DHB. Retrospectively and elegantly, these three crystal structures have underlined the feasibility of fragment merging in drug design. They also corroborate the idea that fragments do not change their binding mode when optimised into leads [44] but their analogues remain precarious prior, see **S5 Fig A and B, Fig 4B and Fig 7A**.

Overall, we have developed a pragmatic ACE fragment screening assay employing the FPPHR method based on the CZE technique. Its low false positive rate arises from the unique set of assay-relevant beneficial properties inherent to CZE. By testing of fewer than 200 compounds, novel promising TTR kinetic stabilisers with nanomolar binding affinity were discovered. Admittedly, this successful FPPHR fragment screening campaign is relatively small (only 129 screened) and easy (binding pockets are deep and “promiscuous” for aromatic compounds). To reaffirm the applicability of the developed FPPHR method in fragment screening, it was deployed again to screen an oncology target in search of novel first-in-class protein-protein interaction inhibitors. 1126 fragments (Maybridge, ThermoFisher) were screened with 47 failed runs, 44 out of the remaining 1079 showed significant Probe Ligand (a short peptide) displacement (4% hit-rate). All hits were subjected to co-crystallisation and/or soaking, dataset of 28 fragments were collected and analysed. Ligand electron density (either full or partial) at the substrate recognition site were observed in all datasets (confidential data). Beyond fragment screening, CZE has been proven useful in answering some fundamental questions scientists faced in early-stage drug discovery. For example, protein analysis of nucleotide bound state (ADP/ATP, GDP/GTP), monitoring compound aqueous solubility or stability over time, and direct measurement of substrate and product levels in an enzyme reaction.

To advocate the use of FPPHR method in FBDD would only be meaningful if there are enough experienced CZE practitioners adopting it. CZE technique proficiency is however heavily reliant on technical knowledge and experiences that might be rare to find in the literature but could be obtained through hands-on practice. It is in hope the published FPPHR example here would stimulate further interests for its use in future FBDD ventures generating highly reliable and quality hits that accelerates the early-stage drug discovery process ultimately benefiting human health.

## Acknowledgements

The research work here formed part of a Ph.D. project (2010-2014) co-supervised by Professor Stephen P. Wood and Dr Carol Austin. All capillary electrophoresis experiments were performed in Selcia Ltd (now part of Eurofins). The author thanks the UCL Wolfson Drug Discovery Unit for the provision of reagents (including recombinant TTR S52P mutant) and use of equipment. Reference normal plasma for ^125^I-T_4_ displacement assay was kindly provided by the Haemophilia Laboratory at the Royal Free Hospital, London, UK. We also gratefully acknowledge access to the synchrotron facilities at ESRF, Grenoble, France and Diamond Light Source, Oxfordshire, United Kingdom.

## Declaration of Conflicting Interests

The author declared no potential conflicts of interest with respect to the research, authorship, and/or publication of this article.

## Support Information

**S1 Fig. Reproducible Analyte Peak Migration Time, Height and Area in Capillary Zone Electrophoresis (CZE) with Superior Separation Efficiency**

Inject Buffer containing 2 μM of transthyretin diluted in Buffer A was prepared. A small volume was injected and separated in a capillary filled with Buffer A. The process was repeated three times with the same capillary and electrophoretic conditions (coated capillary, voltage 30 kV, outlet pressure injection 0.5 psi for 5 seconds). The three electropherograms observed were virtually identical in terms of peak migration time, height and area. This excellent reproducibility and unmatched separation efficiency (even for macromolecules in this case) has made CZE an invaluable technique in fragment screening assays as well as protein analysis.

**S2 Fig. Transthyretin in Complex with Thyroxine T4**

A ribbon representation of transthyretin homotetramer bound to Thyroxine T4. There are two structurally identical thyroxine binding sites in between the dimer-dimer interface with a line of 2-fold symmetry running across the binding channel (red line) . The overall shape of a TTR tetramer resembles of an hourglass and each monomer makes up half of a binding site. Beta-sheet is frequently observed in transthyretin and dissociation of the tetramer can lead to formation of amyloidogenic monomers. Binding of thyroxine or other ligands could stabilise the TTR tetramer by forming non-covalent bonds to each monomer, effectively “securing” them in position.

**S3 Fig. TTR CZE Free 8-ANS Peak Height Restoration by Different Test Compounds**

(**A.**) A fixed volume of Control 2 was injected and separated in a capillary filled with Running Buffer containing different compounds. 500 μM of 2-Aminobenzensulfonamide (Trace A) caused no change in free 8-ANS peak height. 200 μM of 4-Amino-2,6-Dichlorophenol (Trace B) has clearly displaced some 8-ANS from TTR. (**B.**) A fixed volume of Control 1 and Control 2 was injected and separated in a capillary filled with Buffer C twice resulting in identical peaks. Free 8-ANS peak height increased significantly when 250 μM of Diphenylamine was added to the Running Buffer (Trace A), but not with 300 µM of Verapamil (Trace B) *i.e.* a negative. Numeric values atop free 8-ANS peaks indicate migration time (minutes), peak area and height respectively.

**S4 Fig. TTR CZE Free 8-ANS Peak Height Restoration by Different Test Compounds**

(**A.**) A fixed volume of Control 2 was injected and separated in a capillary filled with Buffer C three times (Trace C-E) with highly consistent free 8-ANS peak migration time, area and height. Different compounds were introduced into the Running Buffer. Significant increase in free 8-ANS peak height compared to Control 2 did not happen with 500 µM Acetazolamide (Trace A) but was achieved by Tafamidis at 10 µM (Trace B). Numeric values atop free 8-ANS UV peaks indicate migration time (minutes), peak area and peak height respectively. (**B.**) A sequence of CZE separations was performed with increasing concentrations of 3,5-Dichlorobenzenesulfonamide (SFL000006) added to the Running Buffer. Free 8-ANS UV peak from all traces were superposed and zoomed in omitting the absorbance peak of TTR. Free 8-ANS peak height increased as more SFL000006 was added to the Running Buffer. There was a sharp peak at 3.95 minutes in the blue trace when the compound concentration was at 1 mM. This was treated as a serendipitous spike, possibly due to high compound concentration.

**S5 Fig. TTR Co-Crystal Structure of DIA, DSF and NPA**

(**A.**) The chlorine atoms of 3,5-Dichloroaniline (DIA) are buried deep inside the binding channel and have formed halogen bonds to each of the Ser117 sidechains nearby. (**B.**) A chemical switch from amino to sulphonamide has caused the dichlorobenzene to face the opposite direction towards the binding channel entrance. (**C.**) Unlike DIA and DSF, N-Phenylanthranilic Acid (NPA) occupied both sites in the binding channel. Its diphenylamine is wrapped around by the sidechains of Leu110 and Leu17 whilst the carboxylate forms ionic interaction with Lys15. (**D.**) The crystal structures of DIA (magenta cylinders), NPA (yellow cylinders) and DHB (balls & sticks) in complex with TTR are superposed by Secondary Structure Matching. The corresponding atomic coordinates between these three compounds matched precisely. Numbers near dashed red lines are bond lengths in Angstroms.

**S6 Fig. TTR and SFL001561 Binding**

Binding of SFL001561 to TTR involved a double hydrogen bonding system between the pyrazole ring and the two Ser117 sidechains. This interaction is essential to binding affinity because a methyl substitution on one of the pyrazole nitrogen atoms (SFL001562) had decimated assay activity. The methyl and phenyl substituent contribute to the ligand’s affinity by hydrophobic contact with Leu110 and Leu17 sidechains. There are two possible hydrogen bonds between the amino group and the carbonyl oxygen of Ser117 and Ala108, which is something not seen on previously studied TTR ligands. The binding mode of SFL001561 between the two sites was different, albeit not as obvious as seen for SFL001535. In one site, the phenyl ring is slightly titled along the 2-fold axis compared to the other, another ligand-based structural evidence for binding cooperativity. Numbers near dashed red lines indicate bond length in Angstroms.

**S1 Table. SFL000013 FPPHR Analogue Screen**

**S2 Table. SFL000013 and SFL001535 Analogue Screen**

**S3 Table. SFL001535 Analogue Screen**

**S4 Table. Crystallographic Statistics of Ligand-TTR Co-crystals**

**S1 File. Controls Consistency Record**

**S2 File. Free 8-ANS Peak Height Restoration Titrations from 14 Fragment Hits**

**S3 File. ^125^I-T4 Displacement Curves**

## Notes

https://www.rcsb.org/

### Competing Interest Statement

The authors have declared no competing interest.

