## Supplementary figures and images for "Fragment-Based Drug Discovery for Transthyretin Kinetic Stabilisers Using a Novel Capillary Zone Electrophoresis Method"

### S1 Fig

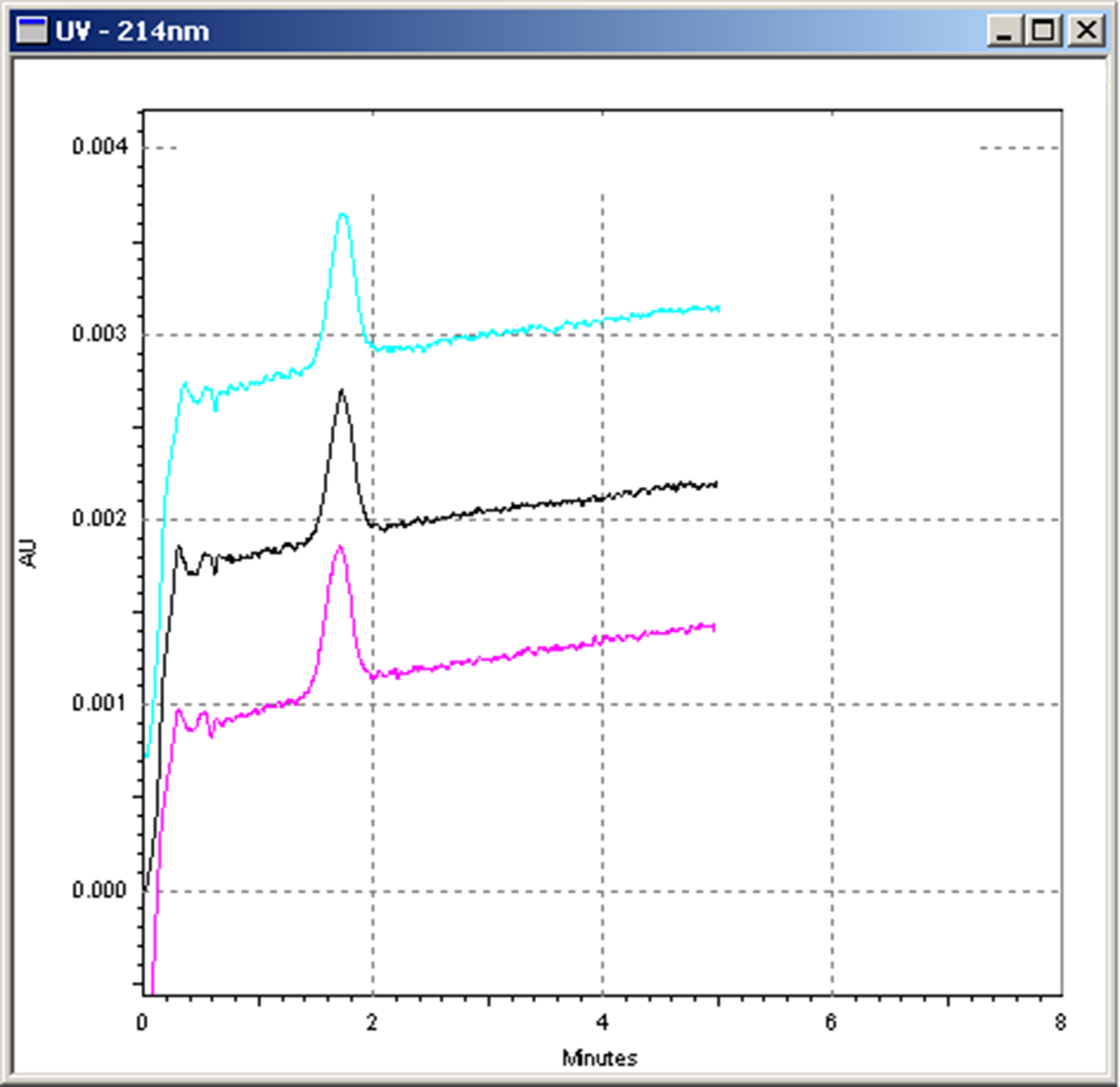

### S2 Fig

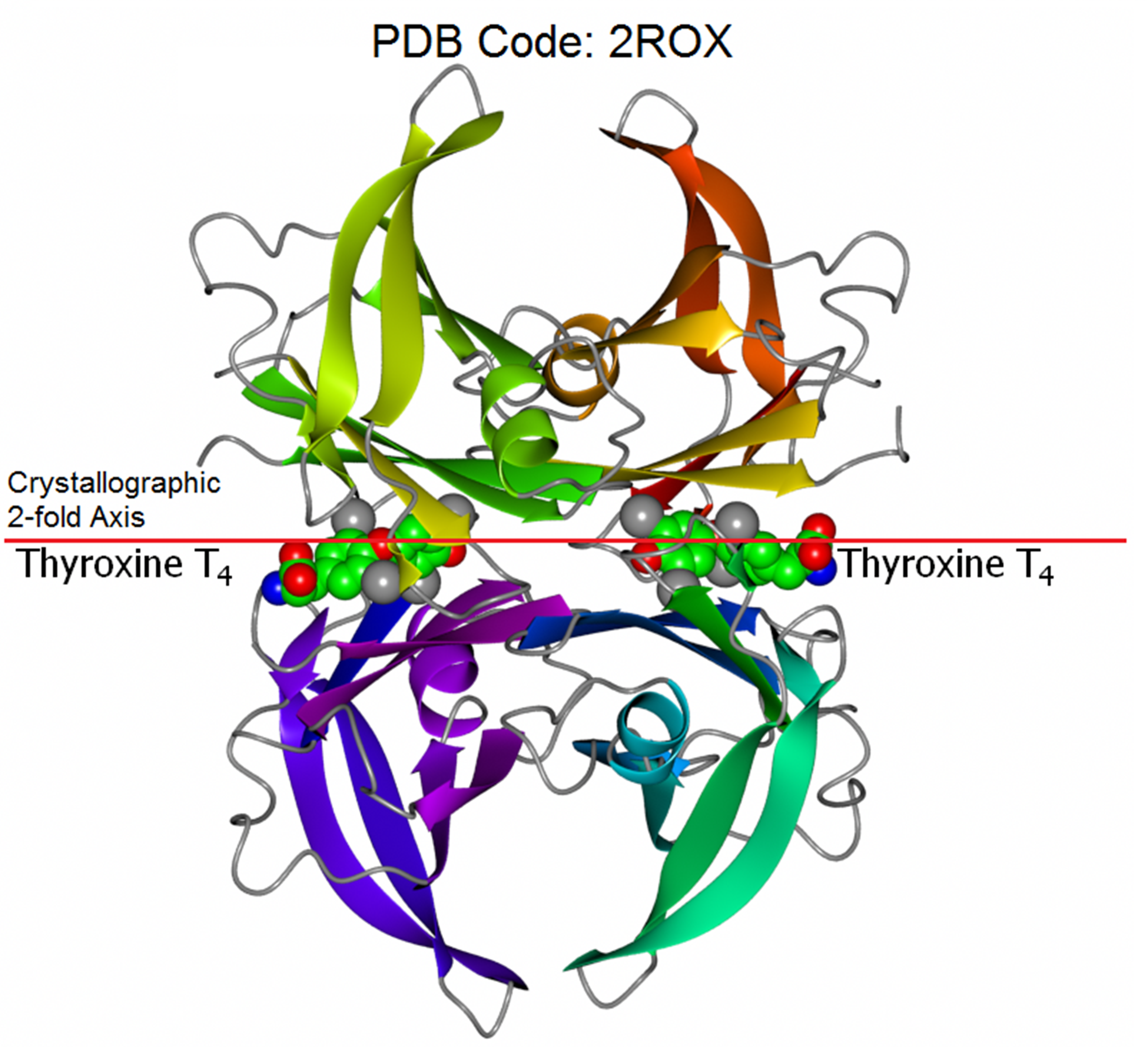

### S3 Fig

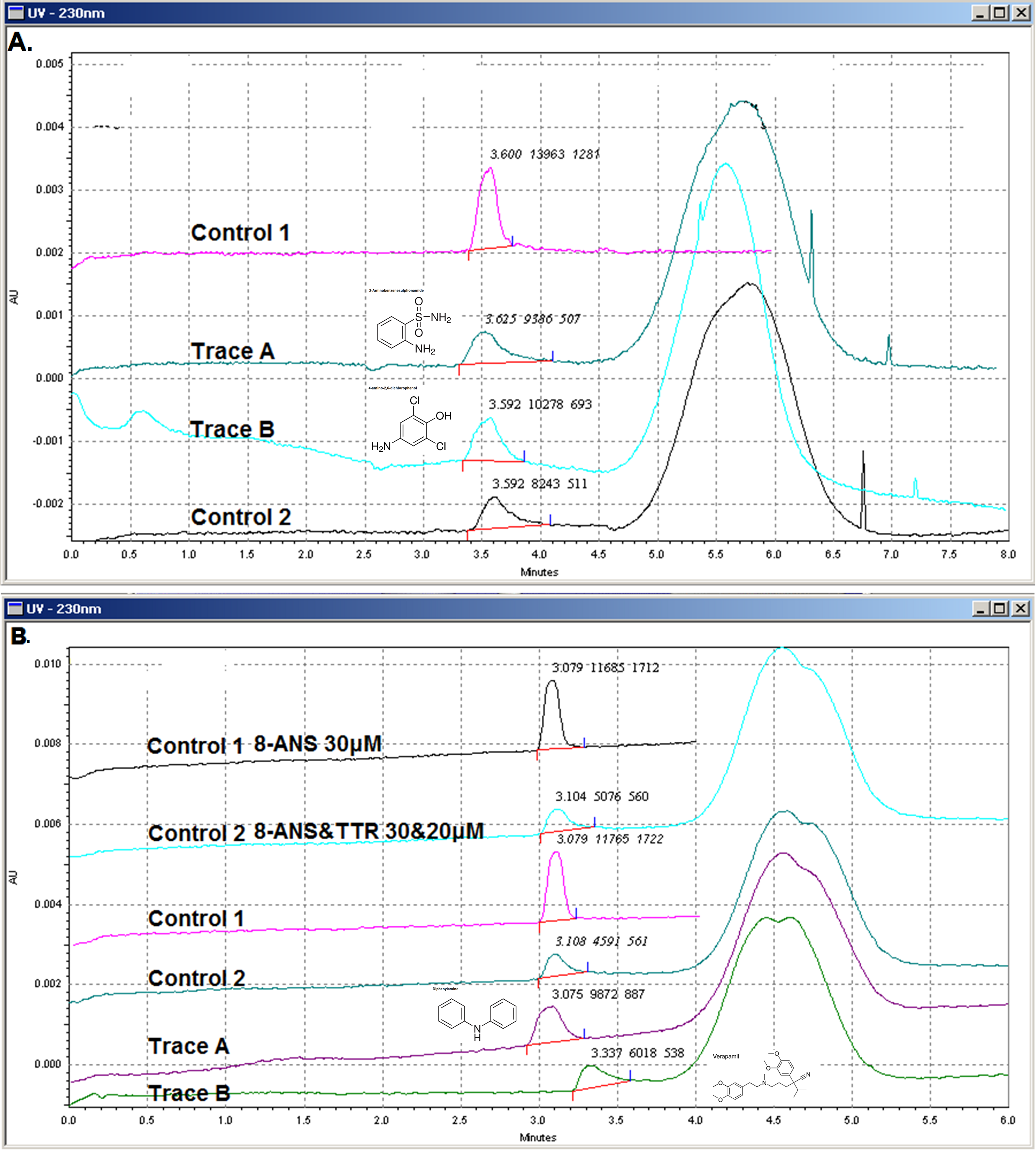

### S4 Fig

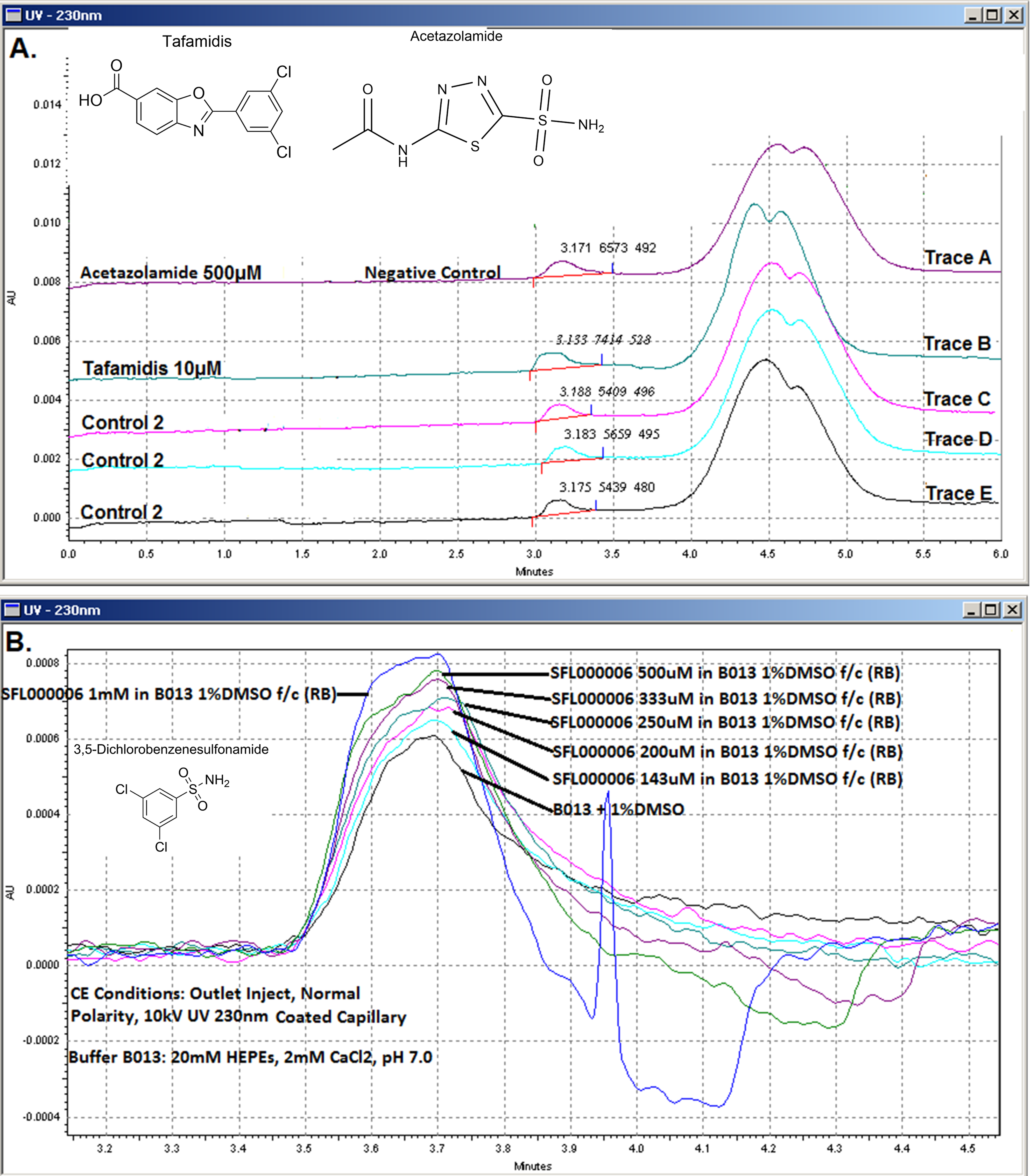

### S6 Fig

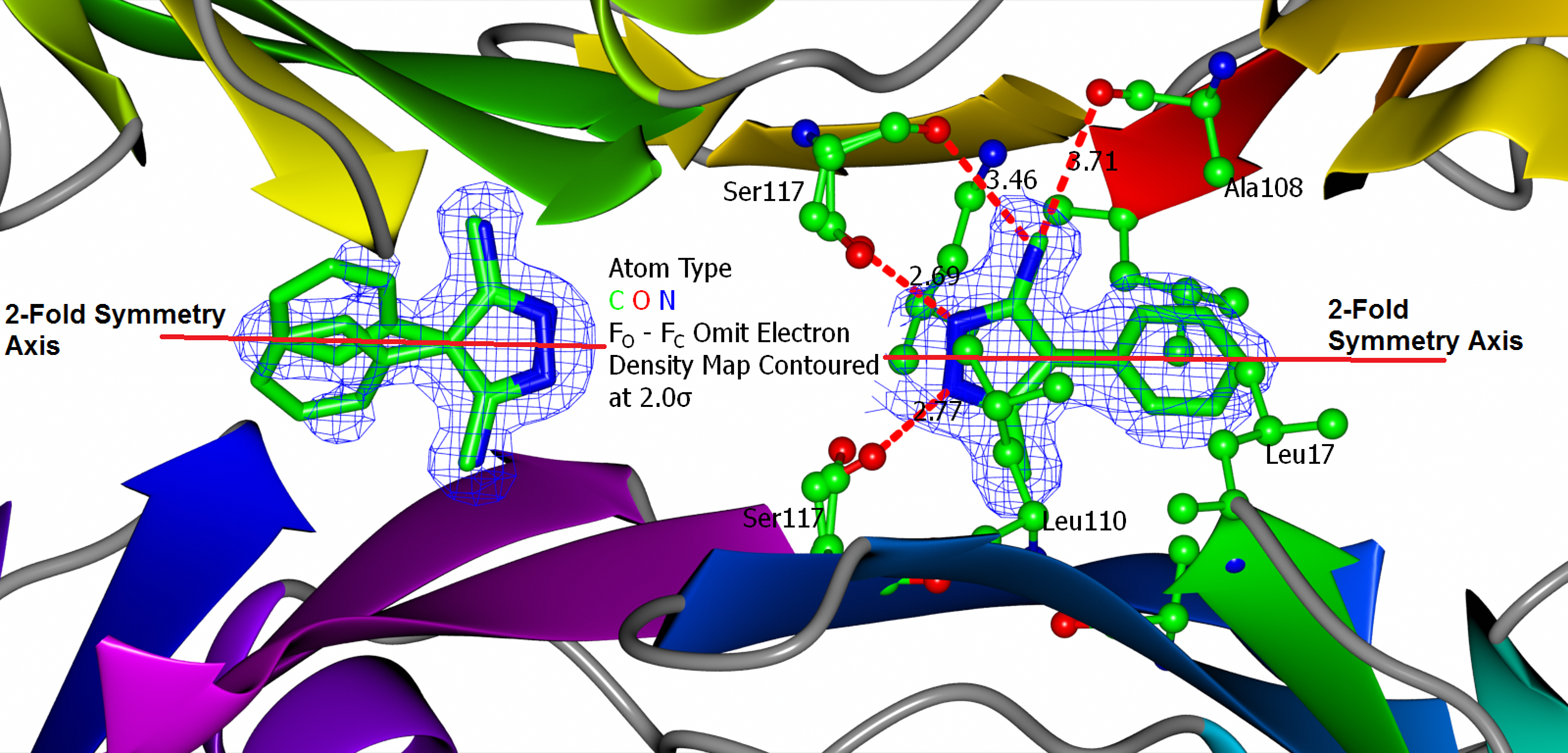
