## Supplementary material for "Fragment-Based Drug Discovery for Transthyretin Kinetic Stabilisers Using a Novel Capillary Zone Electrophoresis Method": S2 File

S2 File. Free 8-ANS Peak Height Restoration Titrations from 14 Fragment Hits

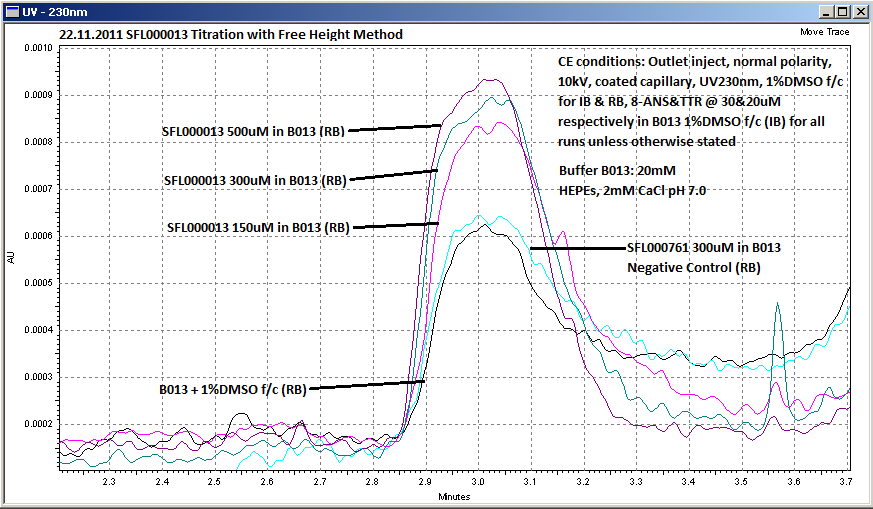

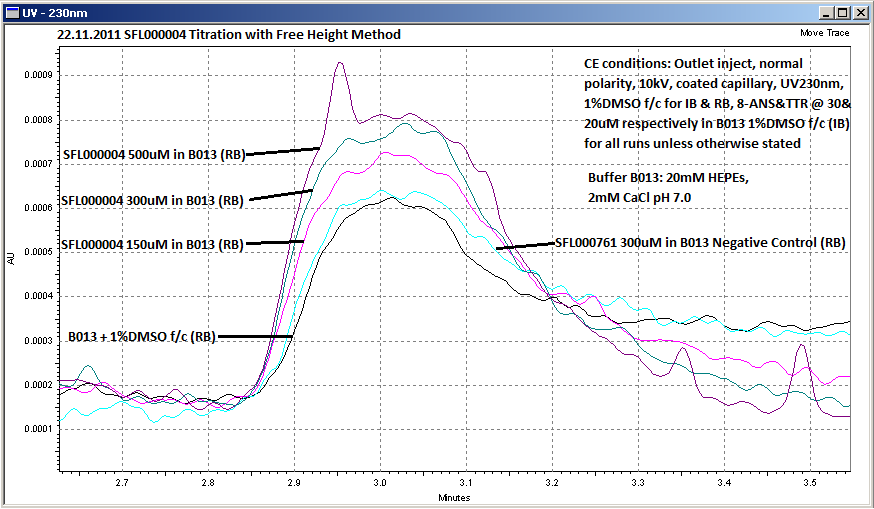

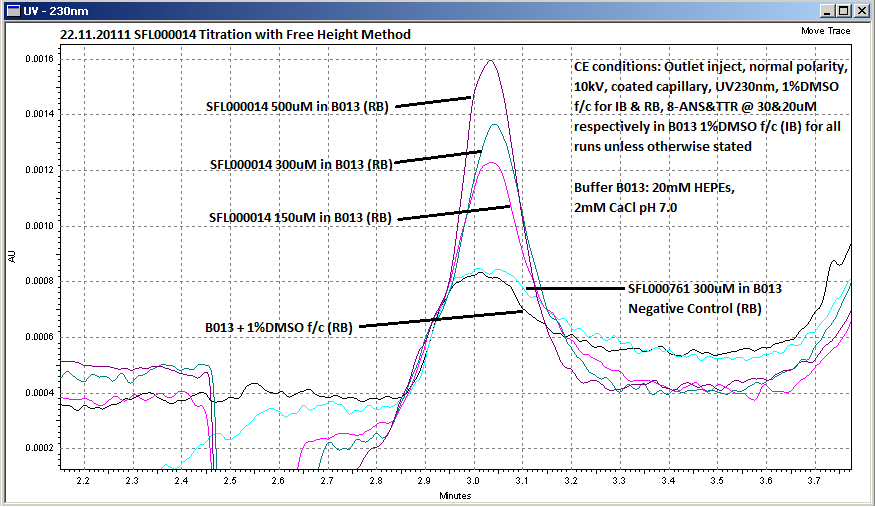

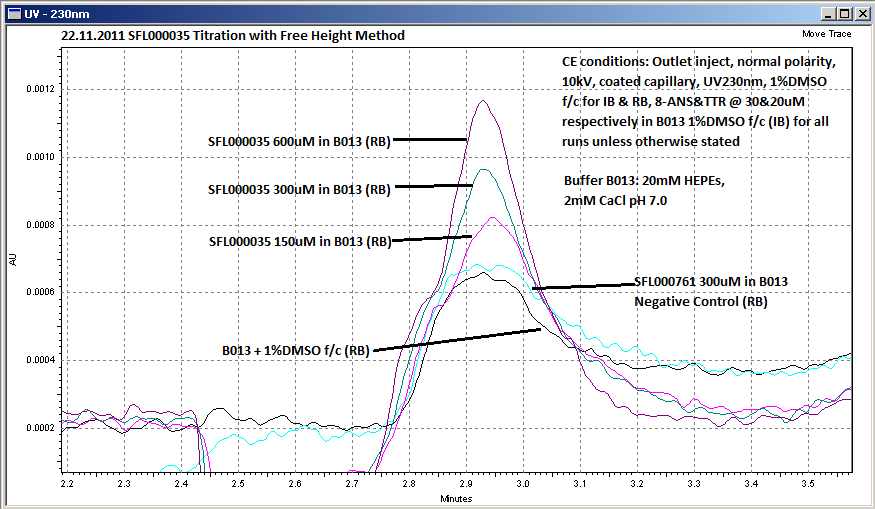

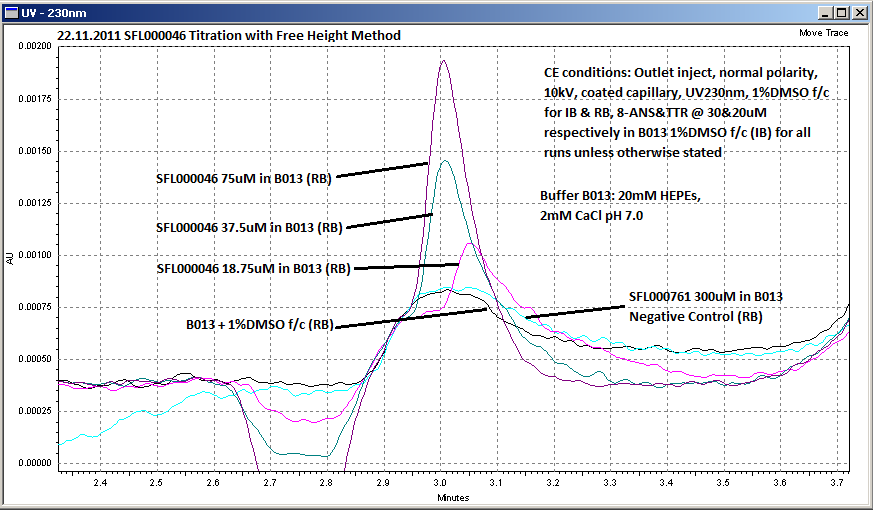

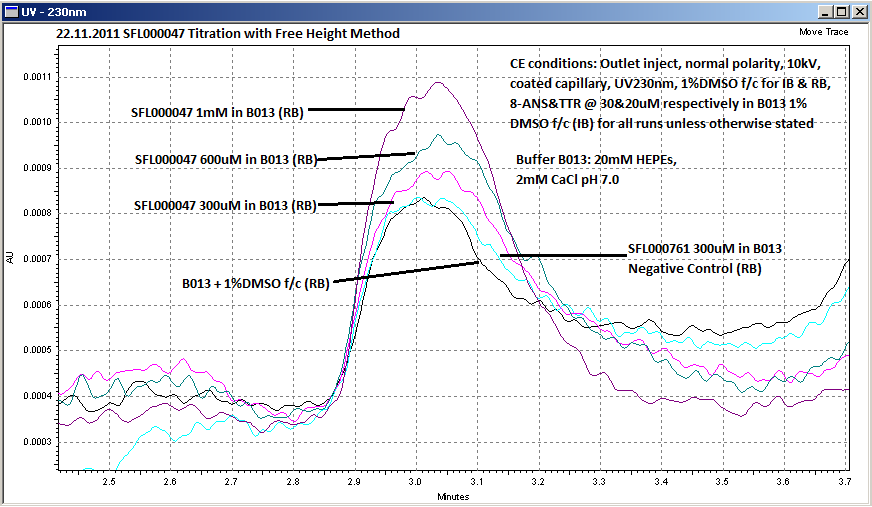

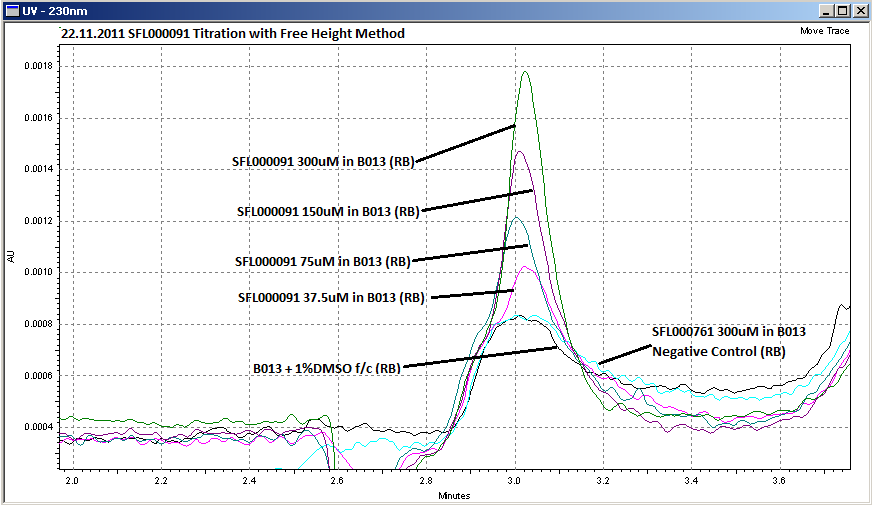

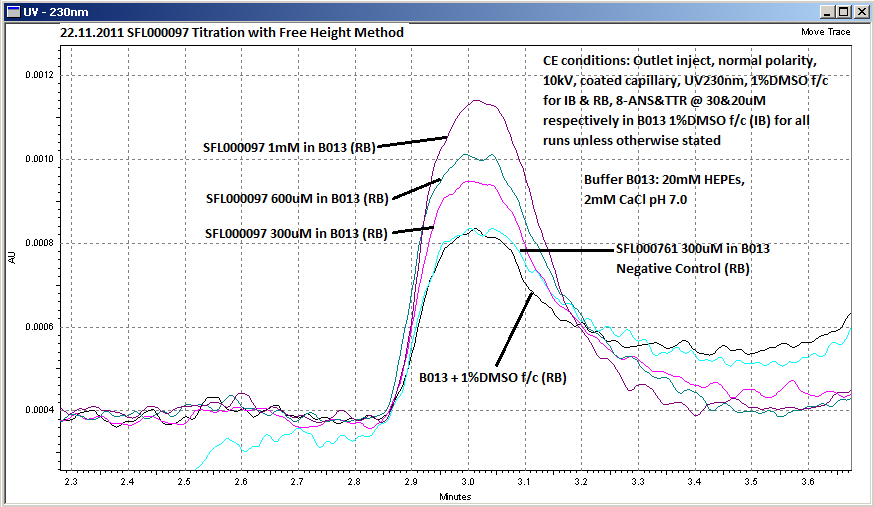

Multiple sequences of CZE separations were performed with increasing concentrations of fragment hits added to the Runing Buffer. Hit titration concentrations ranged from 18.75 µM up to 1mM depending on 8-ANS displacement potency. Free 8-ANS UV peak from all traces were superposed and zoomed in omitting the absorbance peak of TTR.

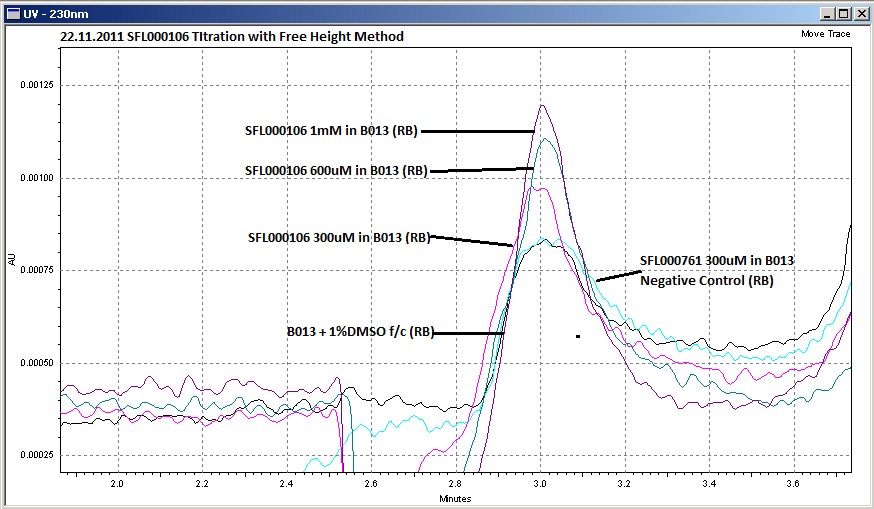

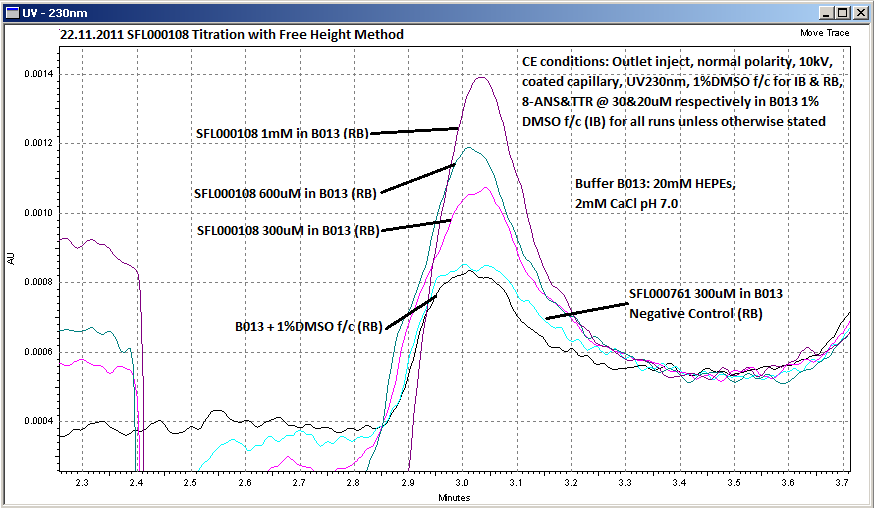

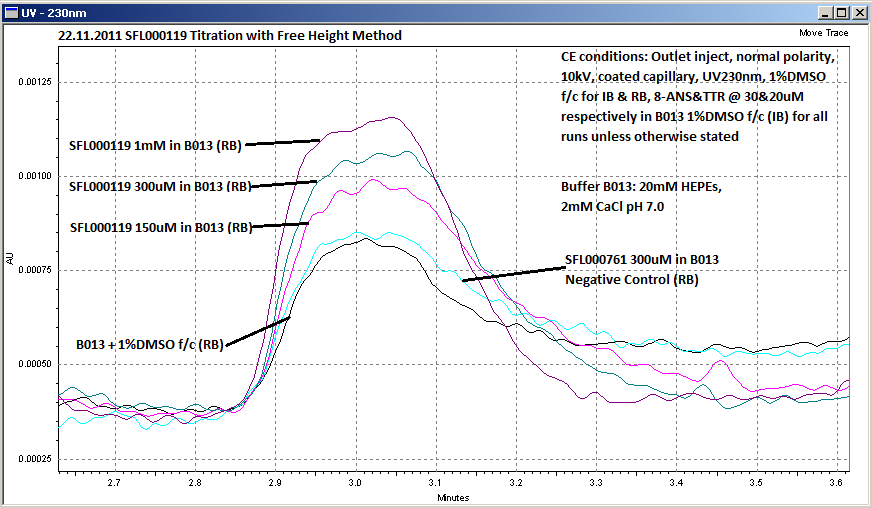

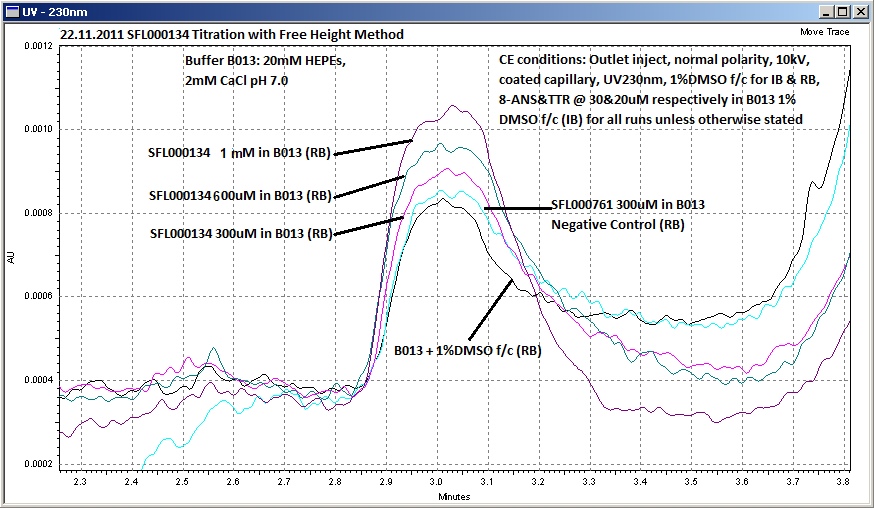

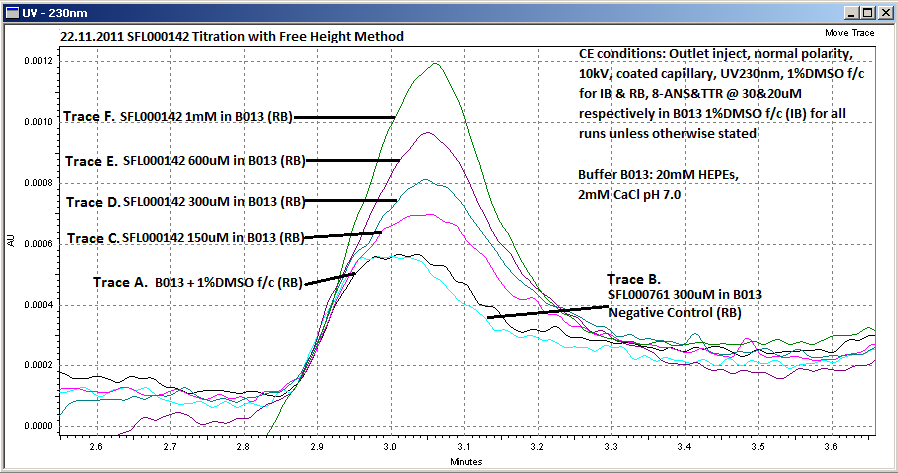

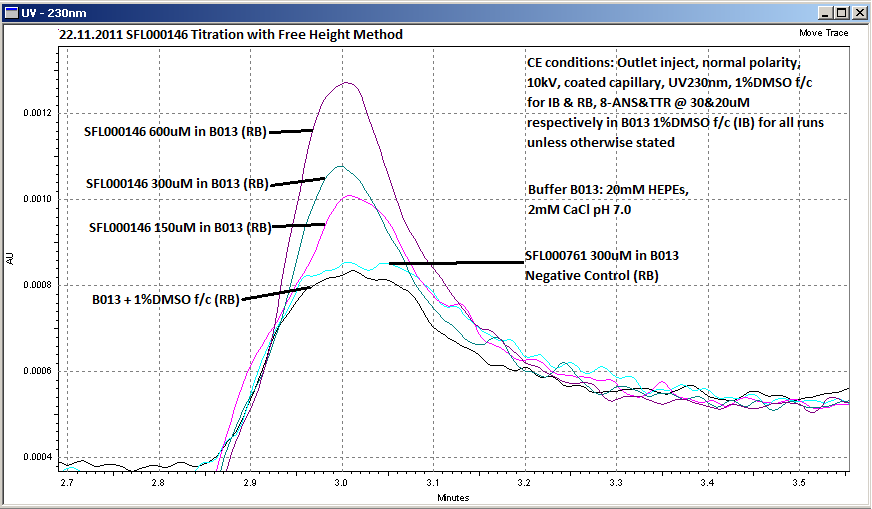
