## Supplementary material for "Fragment-Based Drug Discovery for Transthyretin Kinetic Stabilisers Using a Novel Capillary Zone Electrophoresis Method": S2 Table

S2 Table. SFL000013 and SFL001535 Analogue Screen

| **Fragment ID/Structure** | **Molecular Weight (Number of NHA)** | **FPPHR % at 50 μM ^a^** | **Confirmed by X-ray ^b^** | **Compound Source** |
| --- | --- | --- | --- | --- |
| SFL000013  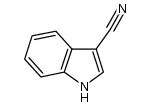 | 142.16 (11) | 18.13% | Yes 9H7G | SFL |
| 218458  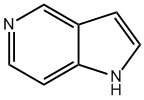 | 118.14 (9) | 5.03% | ND | Selcia Chemical Store |
| A00010944  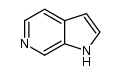 | 118.14 (9) | 3.73% | ND | Selcia Chemical Store |
| 242506  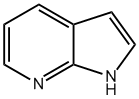 | 118.14 (9) | 7.85% | ND | Selcia Chemical Store |
| A00014954  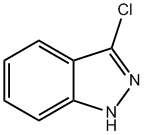 | 152.58 (10) | 24.84% | ND | Selcia Chemical Store |
| 217869  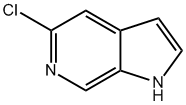 | 153.57 (10) | 12.83% | ND | Selcia Chemical Store |
| A00010562  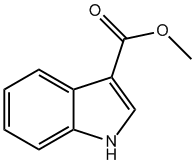 | 175.18 (13) | 16.40% | ND | Selcia Chemical Store |
| SFL001519  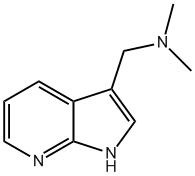 | 175.23 (13) | 1.88% | ND | SFL |
| SFL001561  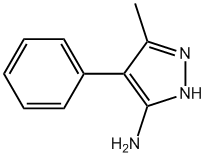 | 173.21 (13) | 34.59% | Yes 9H7J | SFL |
| SFL001562  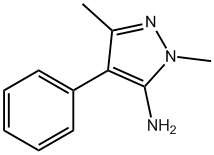 | 187.24 (14) | 7.42% | ND | SFL |
| Z1013760930  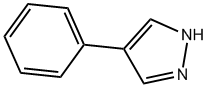 | 144.17 (11) | 14.93% | ND | Selcia Chemical Store |
| OR1969  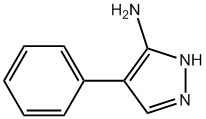 | 159.19 (12) | 13.28% | ND | Selcia Chemical Store |
| 12T-0222  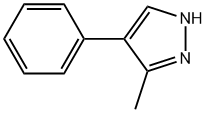 | 158.20 (12) | 25.63% | ND | Selcia Chemical Store |
| 242212  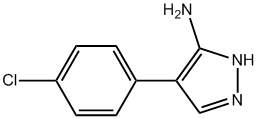 | 193.63 (13) | 25.65% | ND | Selcia Chemical Store |
| SFL001535  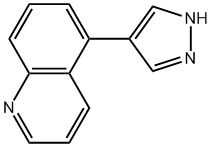 | 195.22 (15) | 60.46% | Yes 9H7I | SFL |
| 94320297  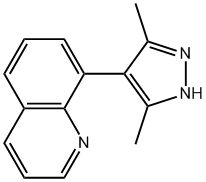 | 223.27 (17) | 59.77% | Yes 9H7K | ChemBridge |
| ^a^ A minimum of 10 % peak height restoration is required to be considered as a binder  ^b^ To be confirmed by X-ray, the fragment in full or part of it had to be observed in the ligand-omitted difference electron density map at a contour level of at least 2σ for one or both binding sites. All structures fitted with ligand are deposited in the Protein Data Bank (PDB) with their PDB ID shown. Crystallographic statistics of all collected datasets can be found in **S4 Table**. ND: Not Done; NHA: Non-Hydrogen Atoms; SFL: Selcia Fragment Library | | | | |
