## Supplementary material for "Fragment-Based Drug Discovery for Transthyretin Kinetic Stabilisers Using a Novel Capillary Zone Electrophoresis Method": S3 File

S3 File. ^125^I-T_4_ Displacement Curves

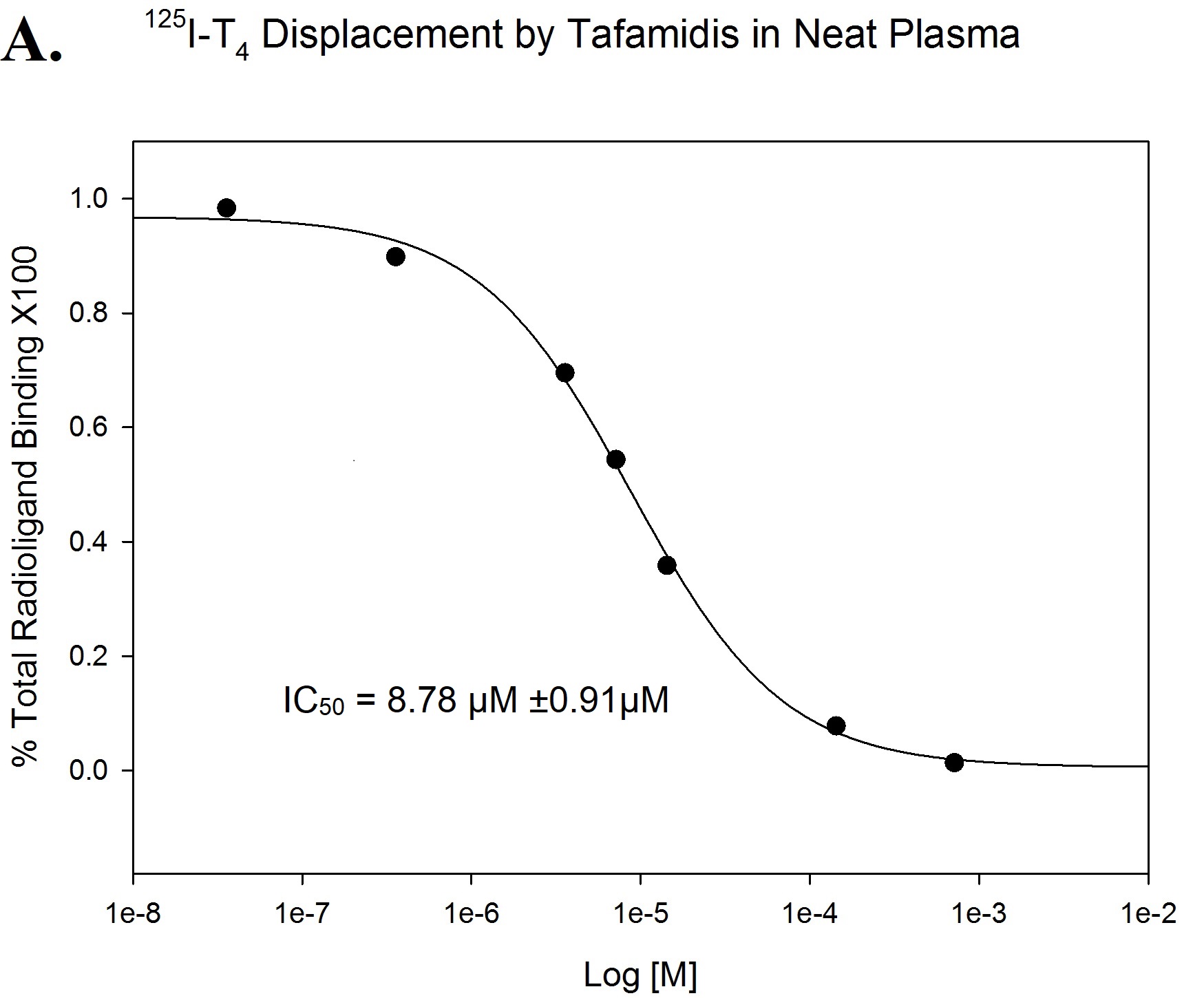

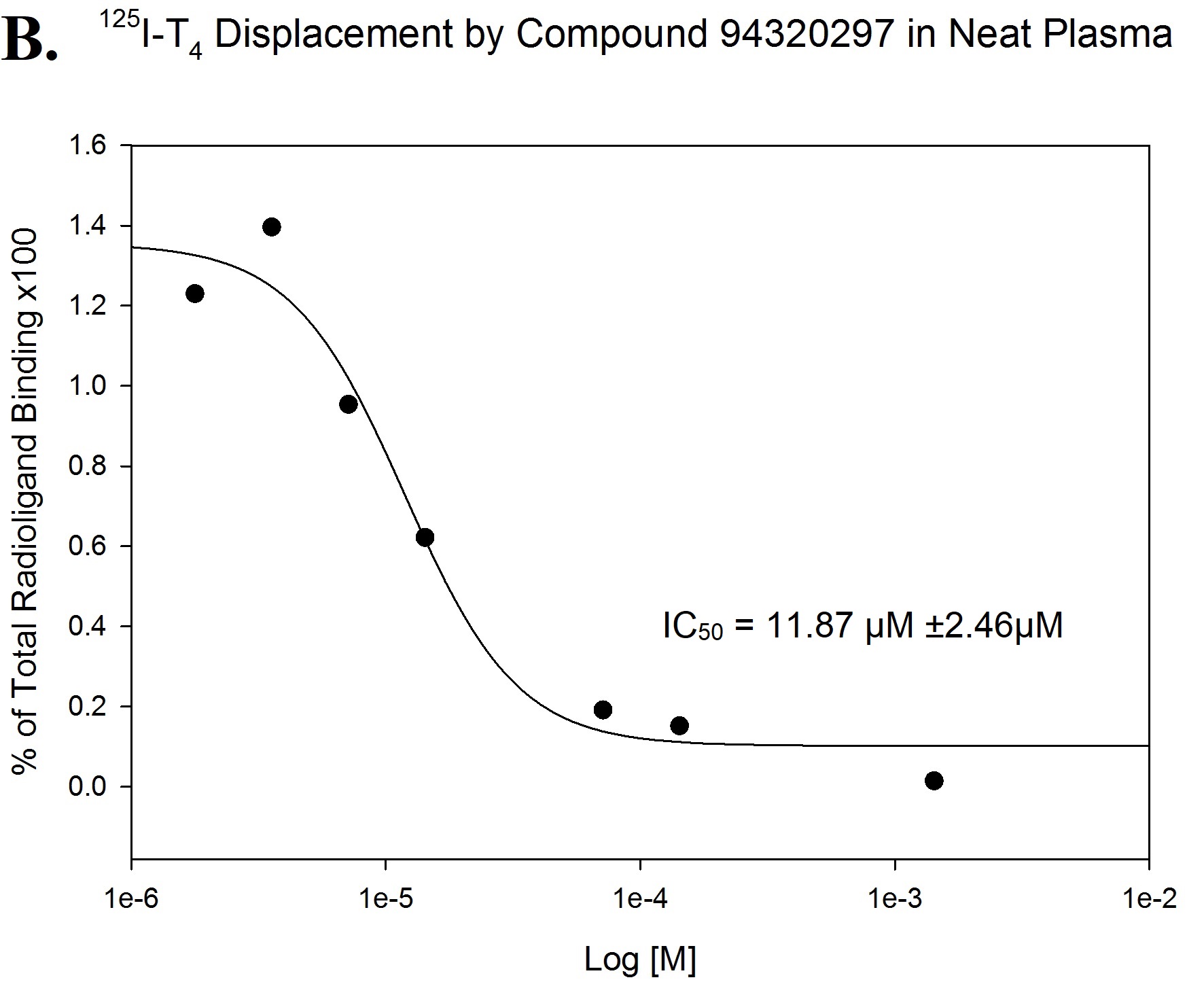

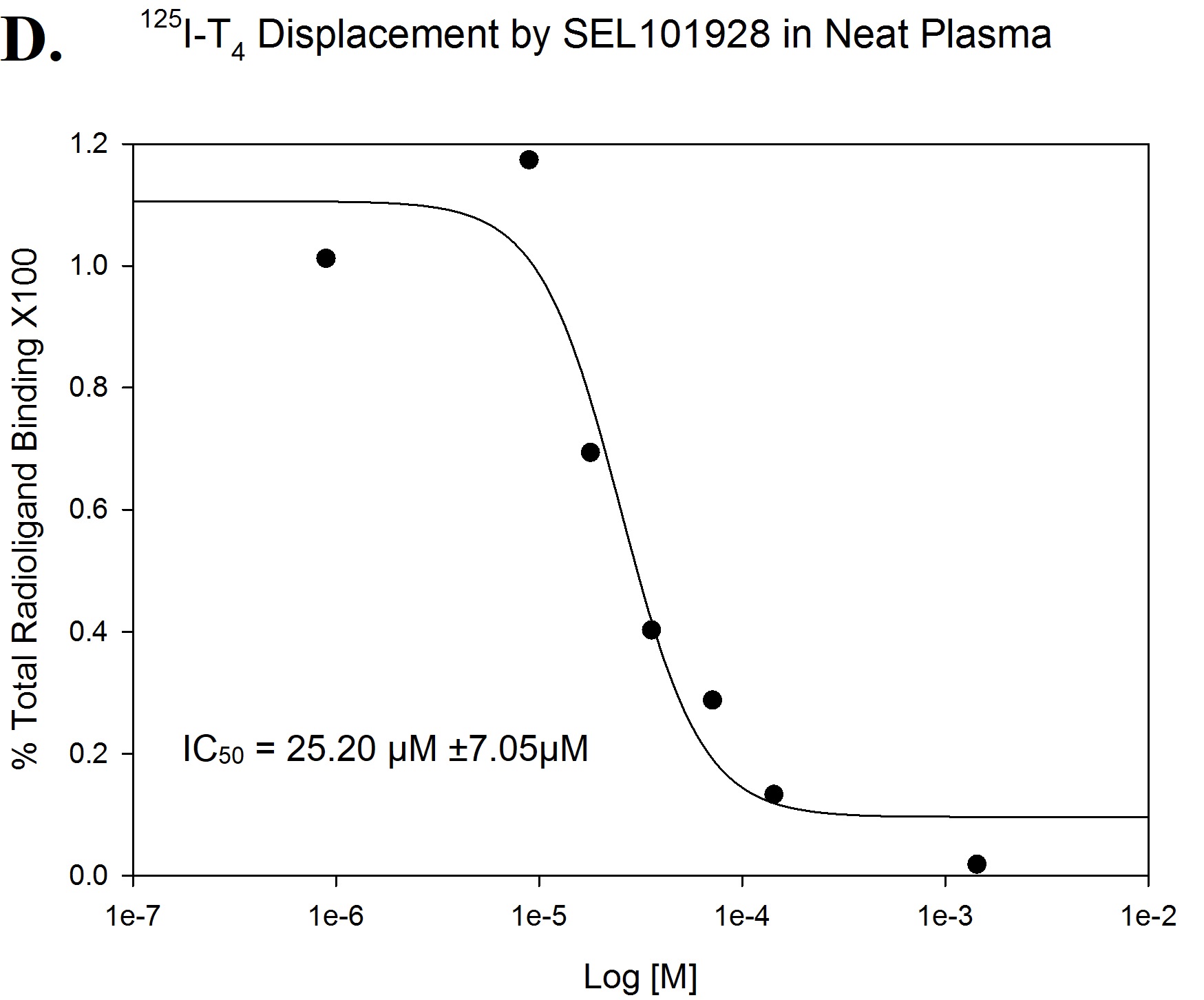

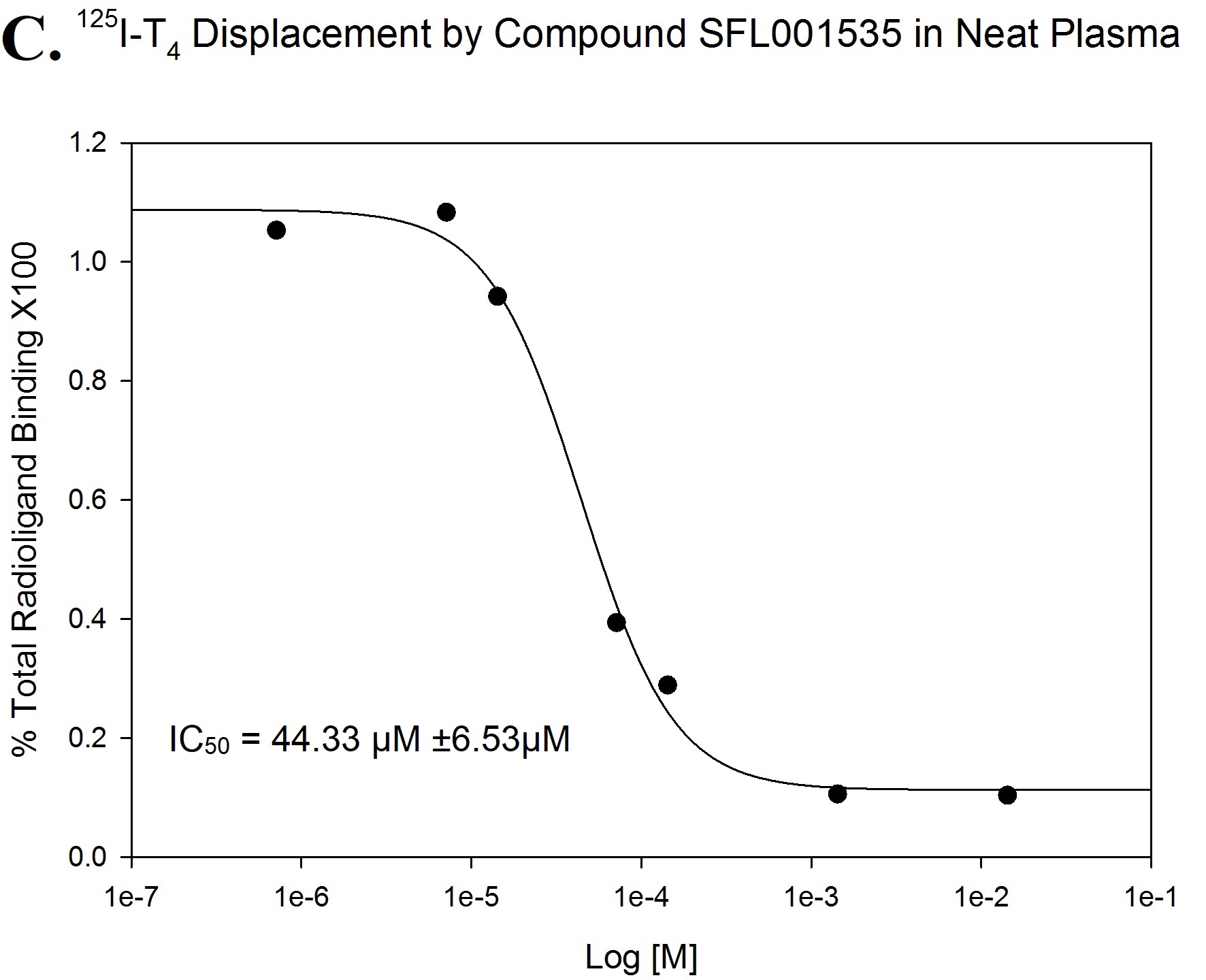

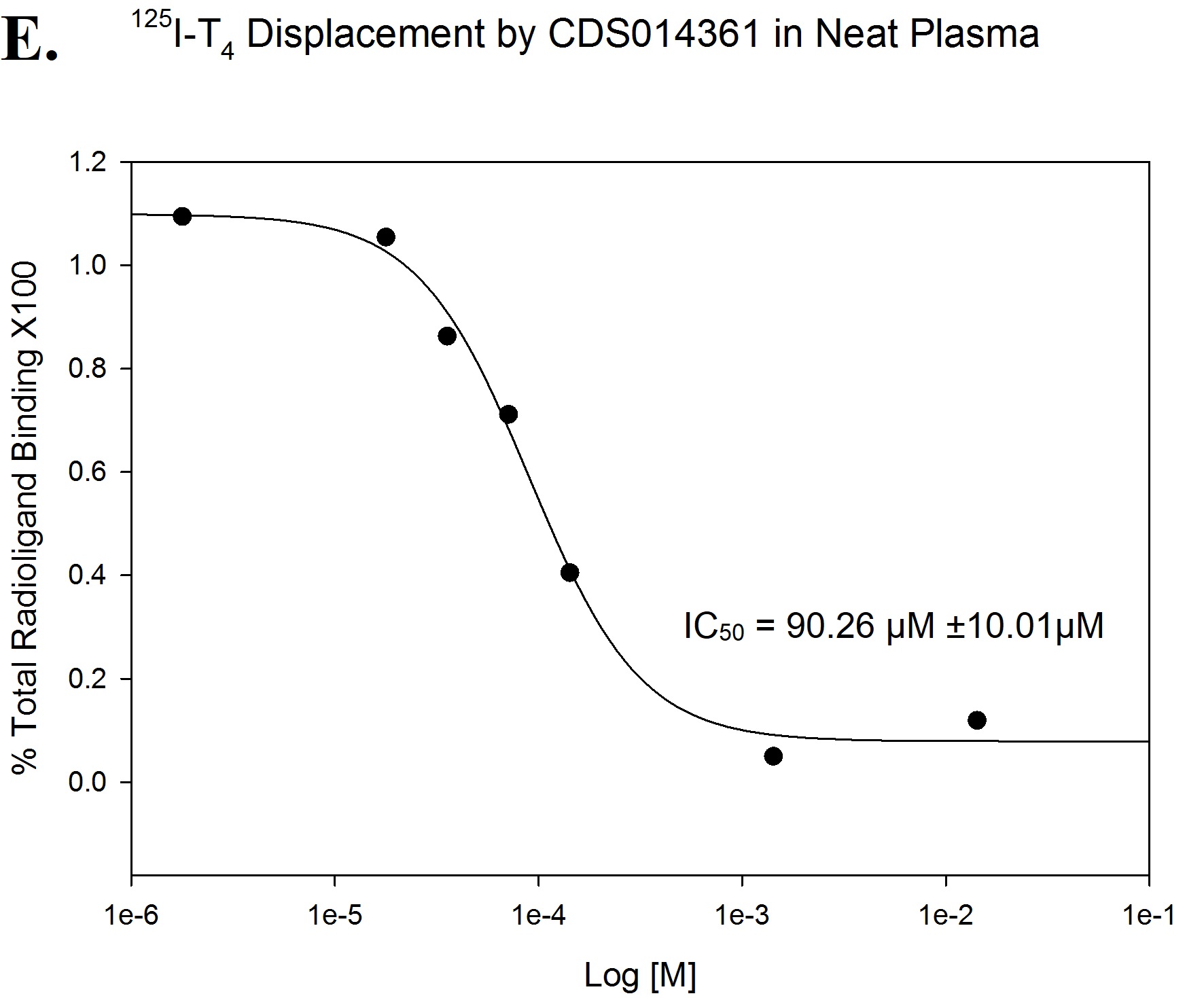

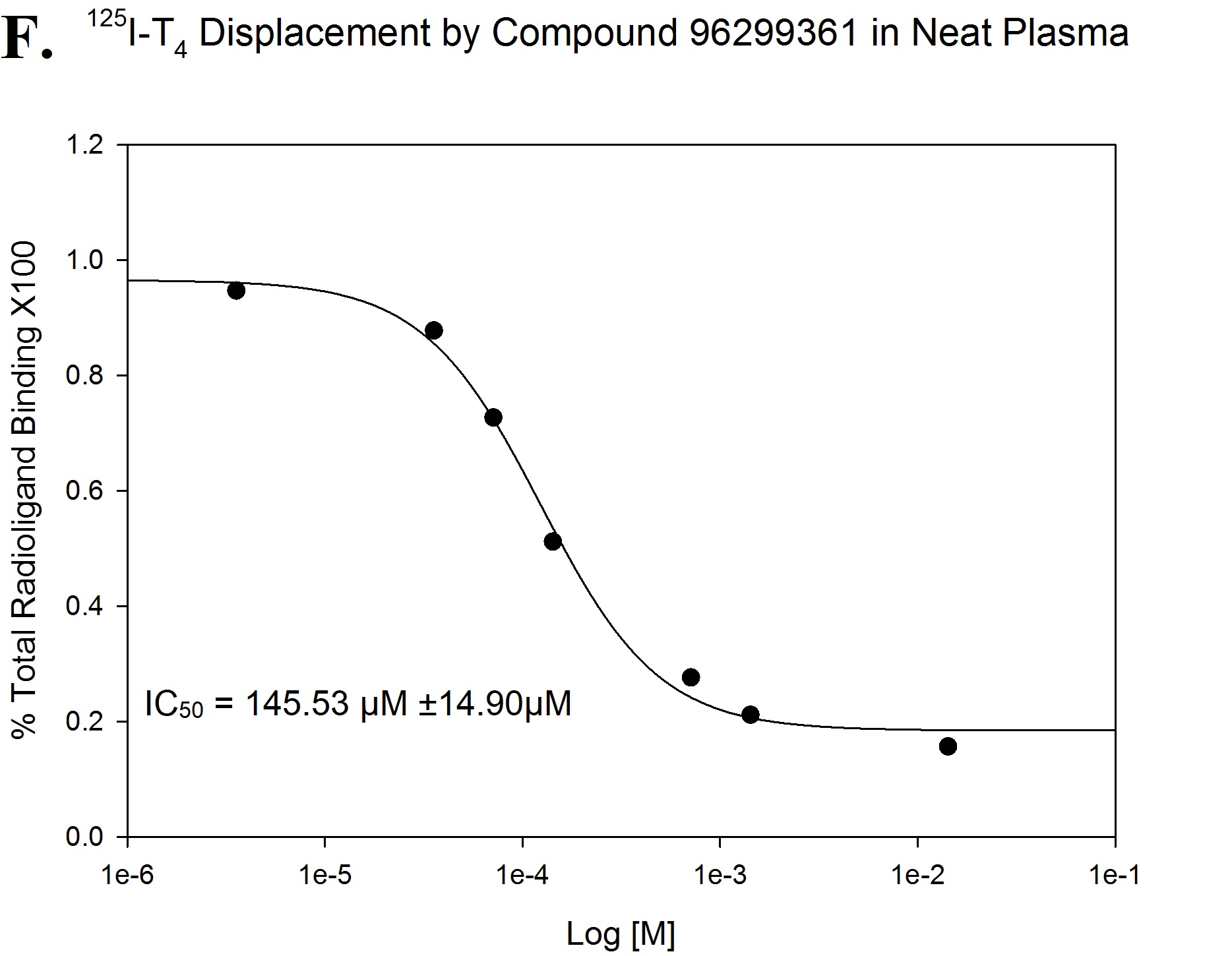

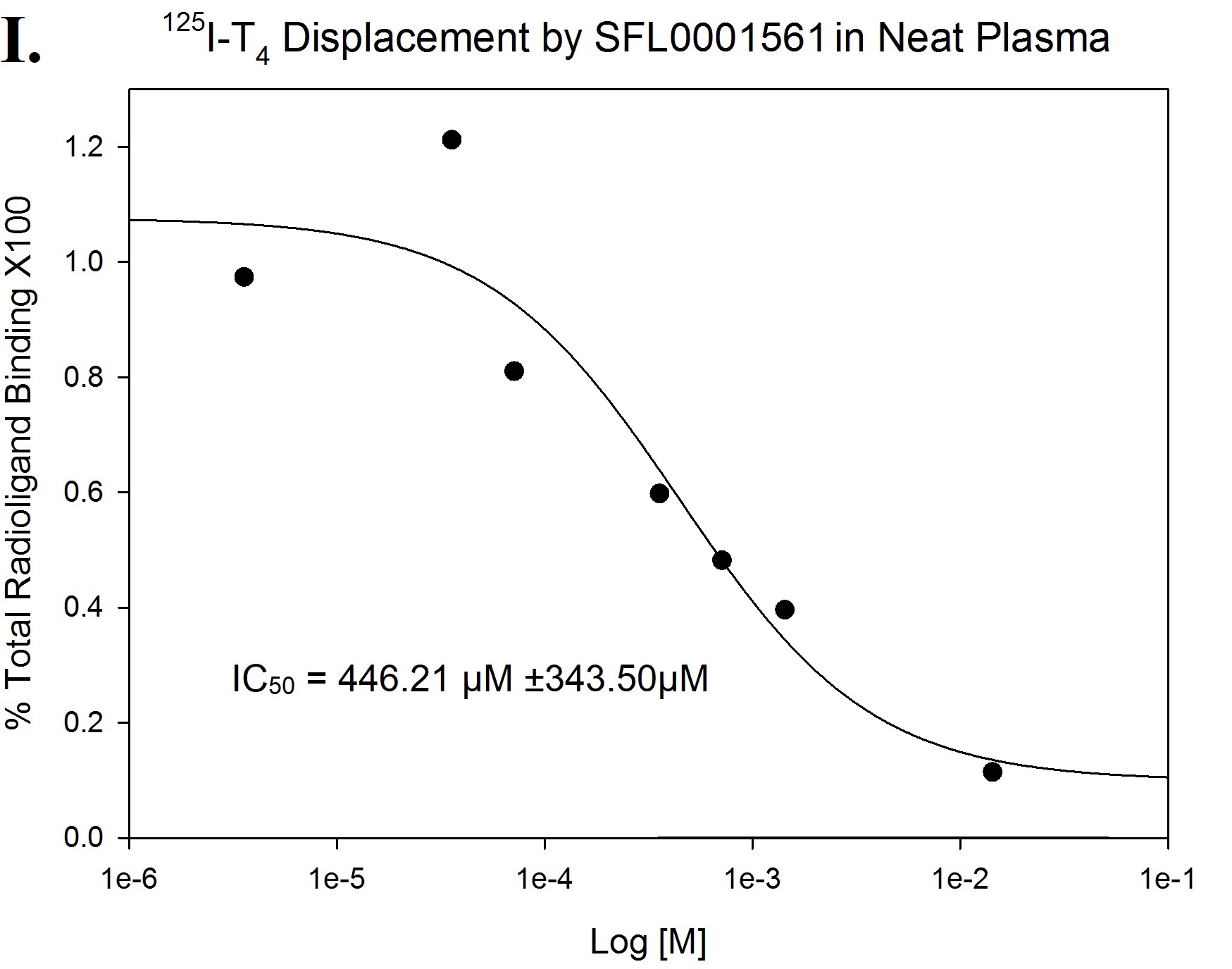

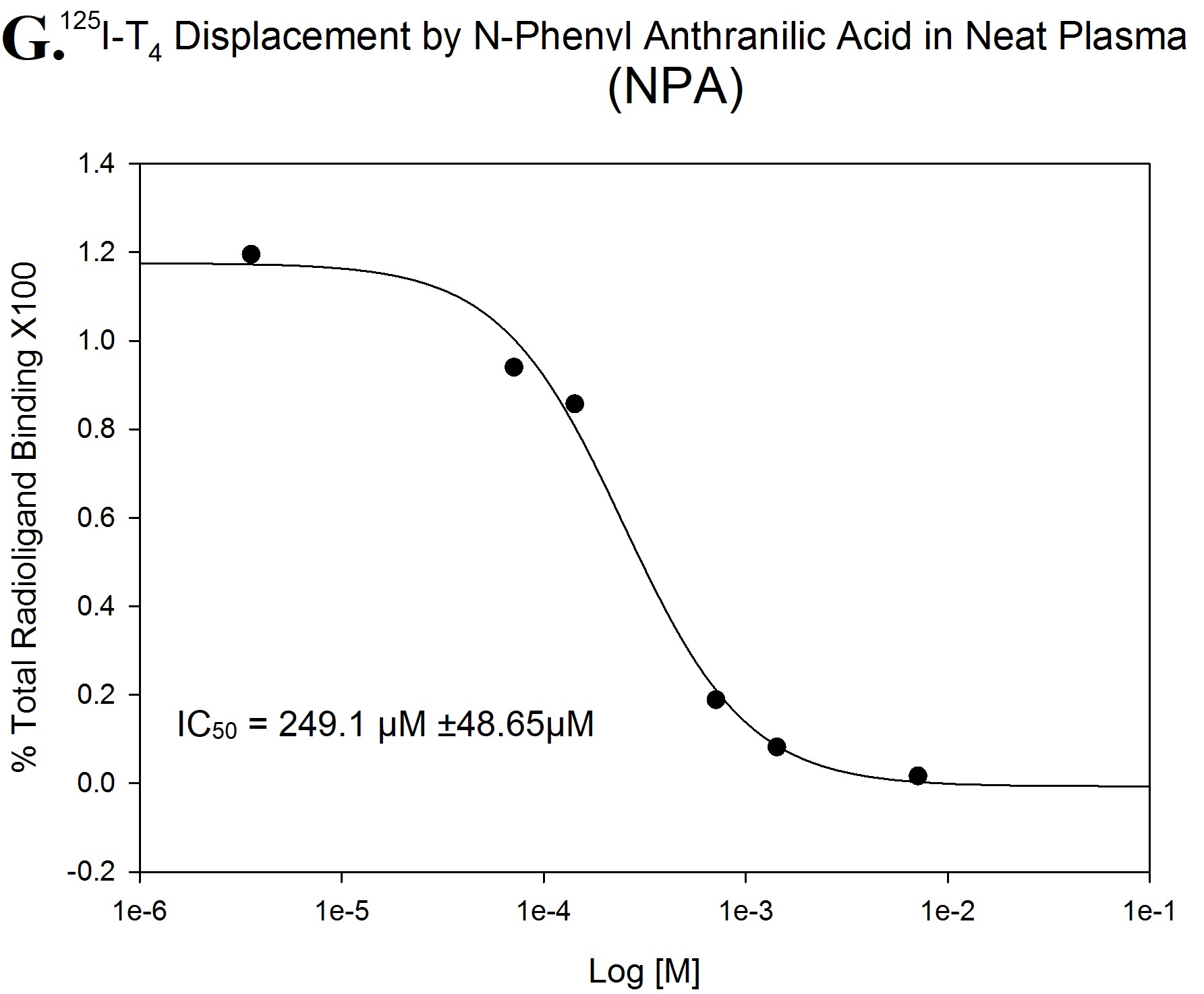

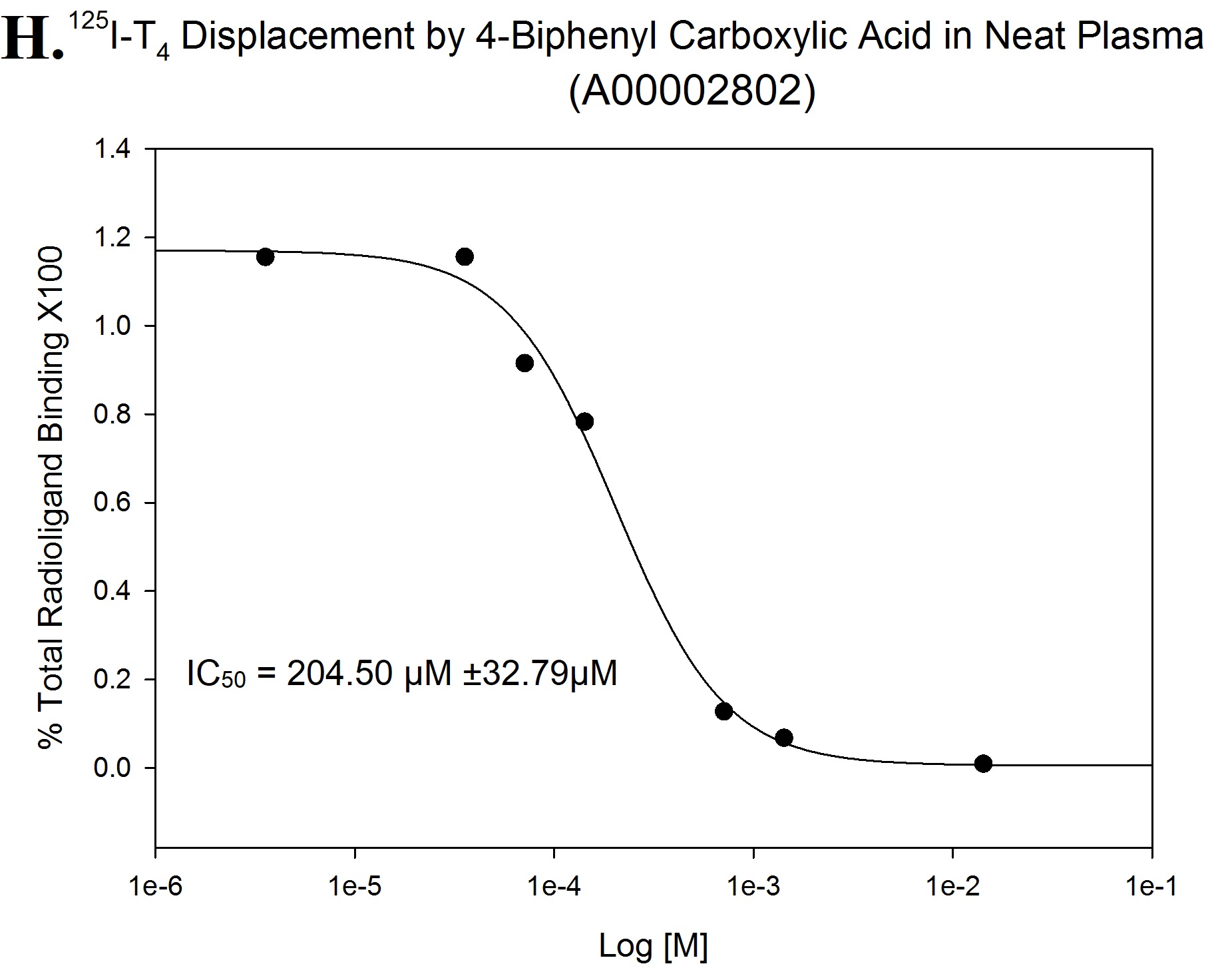

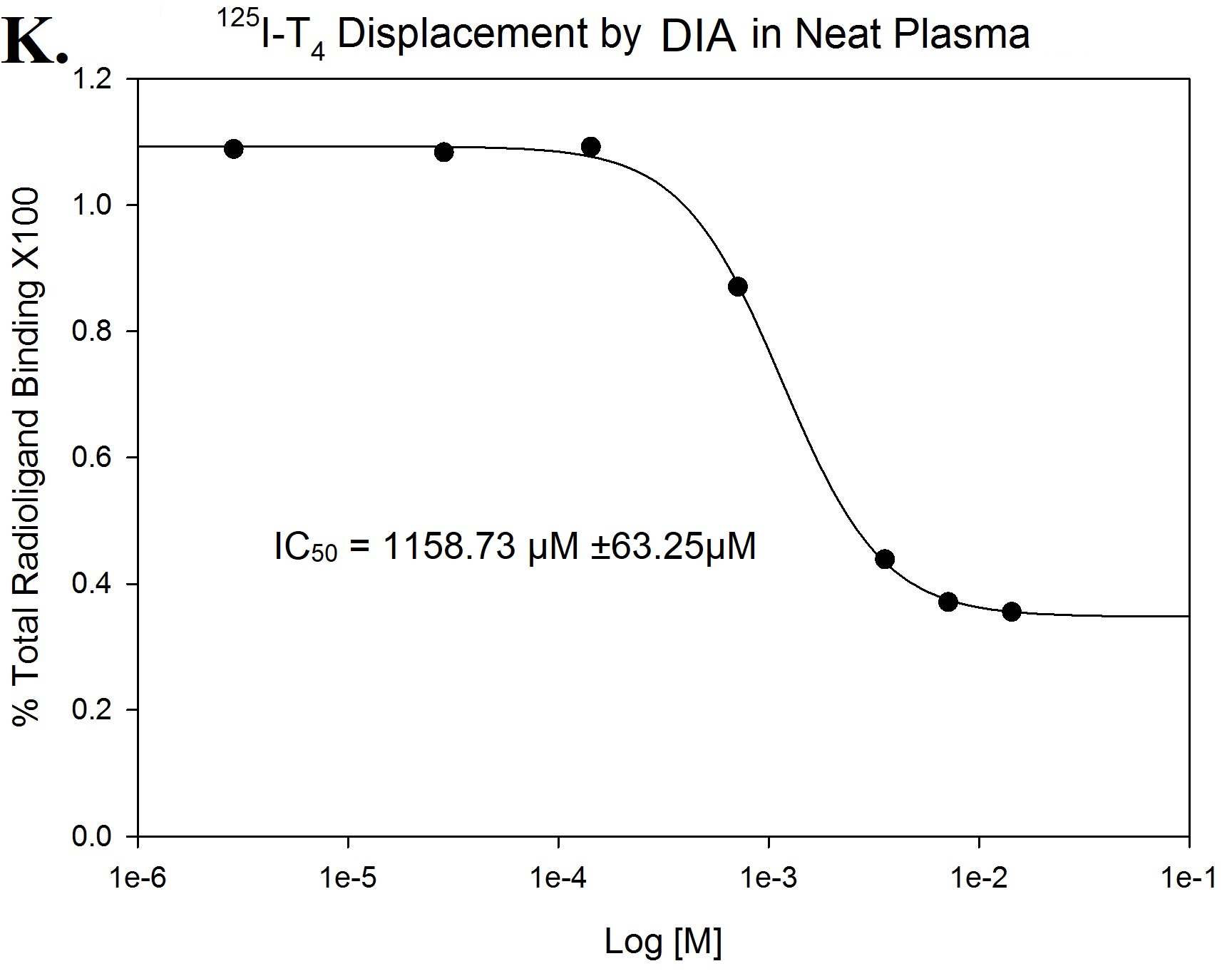

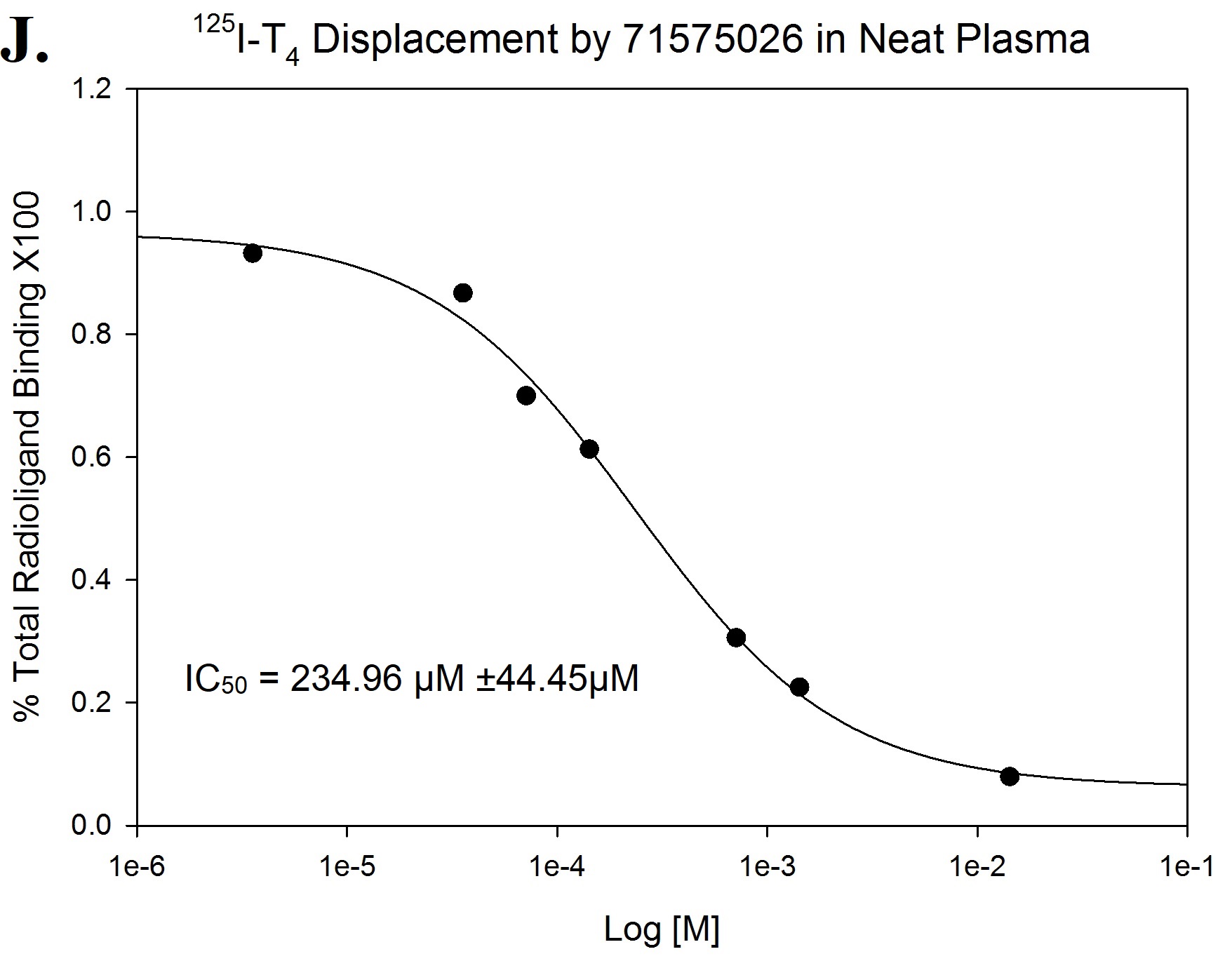

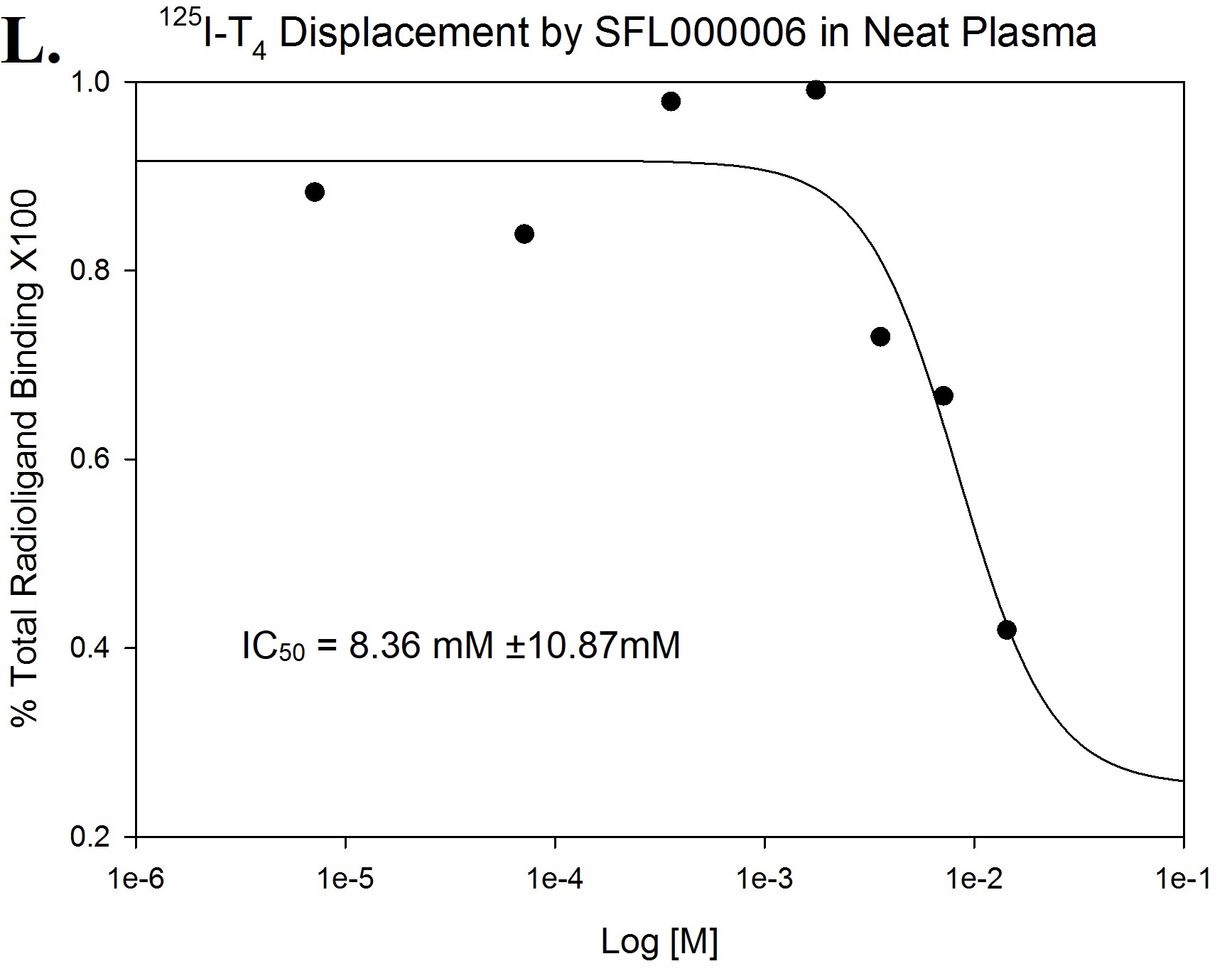

A selection of compounds was tested (3 times) by ^125^I-T_4_ displacement assay in neat plasma and 1 exemplary curve is shown. All were found to be biologically active at different concentrations. 3,5-Dichloroaniline (DIA) plateaued at around 40% total radioligand binding, *i.e.* 60% ^125^I-T_4_ displacement (**K.**). One of the weakest TTR FPPHR fragment hit 3,5-Dichlorobenzenesufonamide (SFL000006) did not reach plateau at maximal concentration (14.3mM) but caused ≈ 60% ^125^I-T_4_ displacement (**L.**). Curves were fitted in SigmaPlot™ (4-Parameter Logistic Function) with calculated IC_50_ and standard error of fitting displayed.
