## Supplementary material for "Fragment-Based Drug Discovery for Transthyretin Kinetic Stabilisers Using a Novel Capillary Zone Electrophoresis Method": S3 Table

S3 Table. SFL001535 Analogue Screen

| **Fragment ID/Structure** | **Molecular Weight (Number of NHA)** | **FPPHR % ^a^** | **Confirmed by X-ray ^b^** | **Compound Source** |
| --- | --- | --- | --- | --- |
| SFL001561  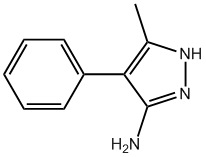 | 173.21 (13) | 34.78% tested at 50 µM | Yes 9H7J | SFL |
| CDS014361  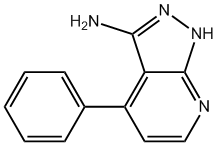 | 210.23 (16) | 36.17% tested at 50 µM | Yes 9H7L | Sigma |
| SFL001535  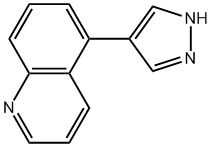 | 195.22 (15) | 34.10% tested at 10 µM | Yes 9H7I | SFL |
| 63633004  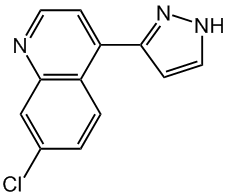 | 229.66 (16) | 22.71% tested at 10 µM | ND | ChemBridge |
| 86471824  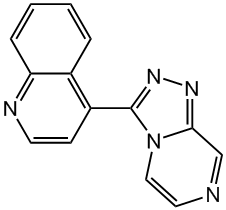 | 247.30 (19) | 10.85% tested at 10 µM | ND | ChemBridge |
| 96299361  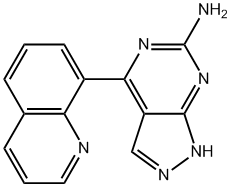 | 262.30 (20) | 23.20% tested at 10 µM | Yes 9H7M | ChemBridge |
| 94320297  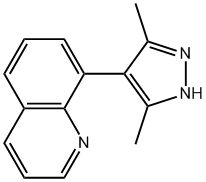 | 223.27 (17) | 33.32% tested at 10 µM | Yes 9H7K | ChemBridge |
| 71575026  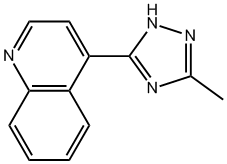 | 210.20 (16) | 17.07% tested at 10 µM | Yes 9H7N | ChemBridge |
| ^a^ A minimum of 10 % peak height restoration is required to be considered as a binder.  ^b^ To be confirmed by X-ray, the fragment in full or part of it had to be observed in the ligand-omitted difference electron density map at a contour level of at least 2σ for one or both binding sites. All structures fitted with ligand are deposited in the Protein Data Bank (PDB) with their PDB ID shown. Crystallographic statistics of all collected datasets can be found in **S4 Table**. ND: Not Done; NHA: Non-Hydrogen Atoms; SFL: Selcia Fragment Library | | | | |
