## Supplementary material for "Fragment-Based Drug Discovery for Transthyretin Kinetic Stabilisers Using a Novel Capillary Zone Electrophoresis Method": S4 Table

S4 Table. Crystallographic Statistics of Ligand-TTR Co-crystals

| **Structure ^a^** | SFL000046 | SFL000091 | SFL000014 | SFL000035 | SFL0000013 | SFL000119 | SFL000004 | SFL000106 |
| --- | --- | --- | --- | --- | --- | --- | --- | --- |
| **Beamline** | DLS I04 | DLS I04-1 | DLS I04 | DLS I04-1 | DLS I04-1 | DLS I04 | ESRF ID29 | DLS I04-1 |
| **Wavelength (Å)** | 0.979 | 0.920 | 0.979 | 0.920 | 0.920 | 0.979 | 0.976 | 0.920 |
| **PDB Code** | 9H7E | Ligand not Fitted | Ligand not Fitted | Ligand not Fitted | 9H7G | Ligand not Fitted | 9H7C | Ligand not Fitted |
| **Ligand-Protein Ratio** | 118:1 | 118:1 | 235:1 | 235:1 | 235:1 | 235:1 | 235:1 | 235:1 |
| **Space Group** | P2_(1)_2_(1)_2_(1)_ | P2_(1)_2_(1)_2_(1)_ | P2_(1)_2_(1)_2 | P2_(1)_2_(1)_2 | P2_(1)_2_(1)_2 | P2_(1)_2_(1)_2 | P2_(1)_2_(1)_2 | P2_(1)_2_(1)_2 |
| **Cell Dimensions (Å)** | 84.12, 41.22, 63.48 | 41.41, 63.38, 83.93 | 83.76, 41.22, 63.66 | 84.14, 41.23, 62.80 | 85.12, 42.18, 64.00 | 84.48, 42.14, 63.46 | 42.03, 85.04, 63.64 | 42.98, 64.80, 85.60 |
| **Resolution (Å)** | 1.43 | 1.55 | 1.40 | 1.48 | 1.49 | 1.36 | 1.27 | 1.43 |
| **Completeness %** | 99.2 | 99.4 | 99.2 | 99.1 | 99.9 | 100 | 98.5 | 99.9 |
| **Multiplicity** | 5.3 | 12.9 | 4.8 | 13.1 | 13.0 | 10.4 | 4.5 | 13.1 |
| **Total Reflections** | 216743 | 417246 | 210609 | 483018 | 499305 | 514348 | 269582 | 588457 |
| **Unique Reflection** | 41021 | 32373 | 43513 | 36865 | 38414 | 514348 | 59970 | 44945 |
| **I/σ^a^** | 10.4 (2.8) | 21.2 (4.0) | 6.4 (2.3) | 25.4 (3.4) | 26.1 (3.6) | 25.2 (4.2) | 13.3 (2.04) | 24.8 (3.9) |
| **R_merge_/R_pim_(I+/-) ^b^** | 0.079 (0.532)/ 0.060 (0.379) | 0.054 (0.667)/ 0.022 (0.268) | 0.100 (0.428)/ 0.068 (0.310) | 0.057 (0.780)/ 0.023 (0.334) | 0.054 (0.711)/ 0.022 (0.306) | 0.045 (0.605)/ 0.021 (0.281) | 0.050 (0.609)/ 0.026 (0.338) | 0.051(0.658)/ 0.021(0.271) |
| **No. of Reflection used for R-free value** | 2059 | 1652 | 2184 | 1841 | 1918 | 2505 | 3038 | 2173 |
| **RMSD of Bond Length(Å)/Angles (°)** | 0.021/2.12 | 0.022/1.91 | 0.020/1.83 | 0.022/1.92 | 0.020/1.965 | 0.022/1.997 | 0.022/2.156 | 0.021/2.050 |
| **R-work/R-free** | 0.237/0.292 | 0.201/0.252 | 0.213/0.263 | 0.190/0.238 | 0.160/0.210 | 0.240/0.270 | 0.166/0.209 | 0.158/0.196 |
| ^a^ all ligands were co-crystallised with wild type TTR unless otherwise stated; ^b^ values in parenthesis are for the highest resolution | | | | | | | | |

| **Structure ^a^** | SFL000108 | SFL000006 | SFL000029 | SFL000097 | SFL001535 | SFL001561 | 94320297 | CDS014361 |
| --- | --- | --- | --- | --- | --- | --- | --- | --- |
| **Beamline** | DLS I04 | DLS I24 | DLS I04-1 | Bruker X8 | DLS I04-1 | DLS I04 | Bruker X8 | ESRF ID29 |
| **Wavelength (Å)** | 0.979 | 0.978 | 0.920 | 1.542 | 0.920 | 0.979 | 1.542 | 0.976 |
| **PDB Code** | Ligand not Fitted | 9H7A | 9H7H | Ligand not Fitted | 9H7I | 9H7J | 9H7K | 9H7L |
| **Ligand-Protein Ratio** | 235:1 | 235:1 | 235:1 | 235:1 | 118:1 | 235:1 | 5:1 | 12:1 |
| **Space Group** | P2_(1)_2_(1)_2 | P2_(1)_2_(1)_2 | P2_(1)_2_(1)_2 | P2_(1)_2_(1)_2 | P2_(1)_2_(1)_2 | P2_(1)_2_(1)_2 | P2_(1)_22_(1)_ | P2_(1)_2_(1)_2 |
| **Cell Dimensions (Å)** | 84.93, 42.06, 63.85 | 85.54, 43.45, 64.63 | 85.640, 42.50, 64.93 | 41.08, 83.64, 62.25 | 85.06, 42.30, 64.77 | 84.48, 42.02, 63.49 | 41.11, 63.07, 83.84 | 42.50, 84.34, 65.74 |
| **Resolution (Å)** | 1.42 | 1.76 | 1.47 | 1.90 | 1.21 | 1.30 | 1.60 | 1.18 |
| **Completeness %** | 99.9 | 99.99 | 99.8 | 99.7 | 98.9 | 100 | 98.6 | 99.3 |
| **Multiplicity** | 10.6 | 12.7 | 12.8 | 5.1 | 8.4 | 10.5 | 4.07 | 5.0 |
| **Total Reflections** | 464784 | 312439 | 525992 | 90160 | 593665 | 590507 | 121129 | 386417 |
| **Unique Reflection** | 43989 | 24574 | 41047 | 17562 | 70465 | 56419 | 29346 | 77826 |
| **I/σ^a^** | 27.9 (3.8) | 18.9 (3.8) | 22.7 (3.6) | 19.62 (7.62) | 19.2 (2.7) | 23.4 (3.4) | 17.78 (4.15) | 13.8 (1.96) |
| **R_merge_/R_pim_(I+/-) ^b^** | 0.041 (0.704)/ 0.019 (0.331) | 0.113 (1.338)/ 0.047 (0.554) | 0.055(0.662)/ 0.023(0.299) | 0.044 (0.182)/ 0.027 (0.105) | 0.050 (0.610)/ 0.026 (0.399) | 0.048 (0.720)/ 0.023 (0.346) | 0.043(0.176)/ 0.022(0.145) | 0.051(0.662)/ 0.025(0.375) |
| **No. of Reflection used for R-free value** | 2212 | 1255 | 2059 | 888 | 3553 | 2855 | 1492 | 3923 |
| **RMSD of Bond Length(Å)/Angles (°)** | 0.022/1.822 | 0.019/1.81 | 0.019/1.819 | 0.019/1.892 | 0.023/2.174 | 0.031/3.028 | 0.019/1.962 | 0.023/2.314 |
| **R-work/R-free** | 0.172/0.217 | 0.196/0.226 | 0.162/0.215 | 0.237/0.281 | 0.154/0.183 | 0.225/0.260 | 0.176/0.235 | 0.171/0.192 |
| ^a^ all ligands were co-crystallised with wild type TTR unless otherwise stated; ^b^ values in parenthesis are for the highest resolution | | | | | | | | |

| **Structure ^a^** | 96299361 | 71575026 | A00002802 | DIA | NPA | 26711816 | SFL001535 (S52P Mutant) |
| --- | --- | --- | --- | --- | --- | --- | --- |
| **Beamline** | DLS I04-1 | ESRF ID29 | DLS I04 | DLS I24 | ESRF ID29 | ESRF ID29 | DLS I04 |
| **Wavelength (Å)** | 0.920 | 0.976 | 0.979 | 0.978 | 0.976 | 0.976 | 0.979 |
| **PDB Code** | 9H7M | 9H7N | 9H7B | 9H7F | 9H7D | 9H7O | 9H7P |
| **Ligand-Protein Ratio** | 23:1 | 12:1 | 7:1 | 235:1 | 117:1 | 23:1 | 235:1 |
| **Space Group** | P2_(1)_2_(1)_2 | P2_(1)_2_(1)_2 | P2_(1)_2_(1)_2 | P2_(1)_2_(1)_2 | P2_(1)_2_(1)_2 | P2_(1)_2_(1)_2 | P2(1)2(1)2 |
| **Cell Dimensions (Å)** | 85.03, 42.13, 63.73 | 42.35, 84.66, 65.39 | 83.49, 41.10, 62.77 | 85.11, 43.80, 64.76 | 41.15, 84.08, 62.30 | 42.07.15, 85.04, 64.27 | 84.86, 41.90, 63.59 |
| **Resolution (Å)** | 1.27 | 1.26 | 1.51 | 1.24 | 1.23 | 1.24 | 1.36 |
| **Completeness %** | 99.1 | 97.5 | 99.8 | 99.91 | 97.5 | 96.7 | 99.6 |
| **Multiplicity** | 13.1 | 4.1 | 4.8 | 12.7 | 4.8 | 4.6 | 4.6 |
| **Total Reflections** | 792870 | 259495 | 165005 | 901013 | 299244 | 290801 | 225550 |
| **Unique Reflection** | 60596 | 62554 | 34677 | 71035 | 61779 | 63682 | 49469 |
| **I/σ^a^** | 22.5 (3.9) | 13.8 (1.81) | 12.5 (2.4) | 28.8 (3.7) | 11.04 (2.17) | 14.07 (1.83) | 10.9 (2.1) |
| **R_merge_/R_pim_(I+/-) ^b^** | 0.051(0.640)/ 0.021(0.265) | 0.049(0.610)/ 0.026(0.399) | 0.056 (0.491)/ 0.043 (0.375) | 0.079 (0.777)/ 0.033 (0.344) | 0.061(0.342)/ 0.030(0.167) | 0.045(0.592)/ 0.023(0.327) | 0.055 (0.525) /0.041(0.402) |
| **No. of Reflection used for R-free value** | 3059 | 3169 | 1743 | 3507 | 3112 | 3208 | 2503 |
| **RMSD of Bond Length(Å)/Angles (°)** | 0.026/2.316 | 0.023/2.008 | 0.020/1.932 | 0.022/2.07 | 0.025/2.31 | 0.025/1.98 | 0.022/2.111 |
| **R-work/R-free** | 0.150/0.191 | 0.164/0.205 | 0.169/0.228 | 0.156/0.178 | 0.162/0.188 | 0.159/0.193 | 0.165/0.205 |
| ^a^ all ligands were co-crystallised with wild type TTR unless otherwise stated; ^b^ values in parenthesis are for the highest resolution | | | | | | | |
