## Supplementary material for "Fragment-Based Drug Discovery for Transthyretin Kinetic Stabilisers Using a Novel Capillary Zone Electrophoresis Method": S5 Fig

S5 Fig. TTR Co-Crystal Structure of DIA, DSF and NPA

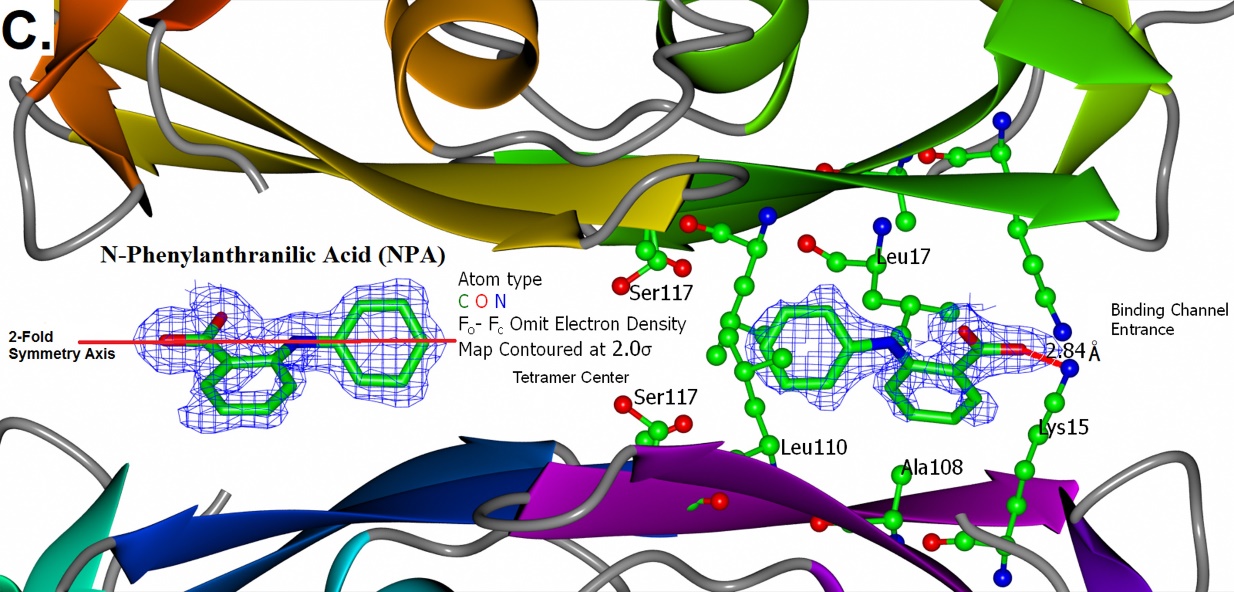

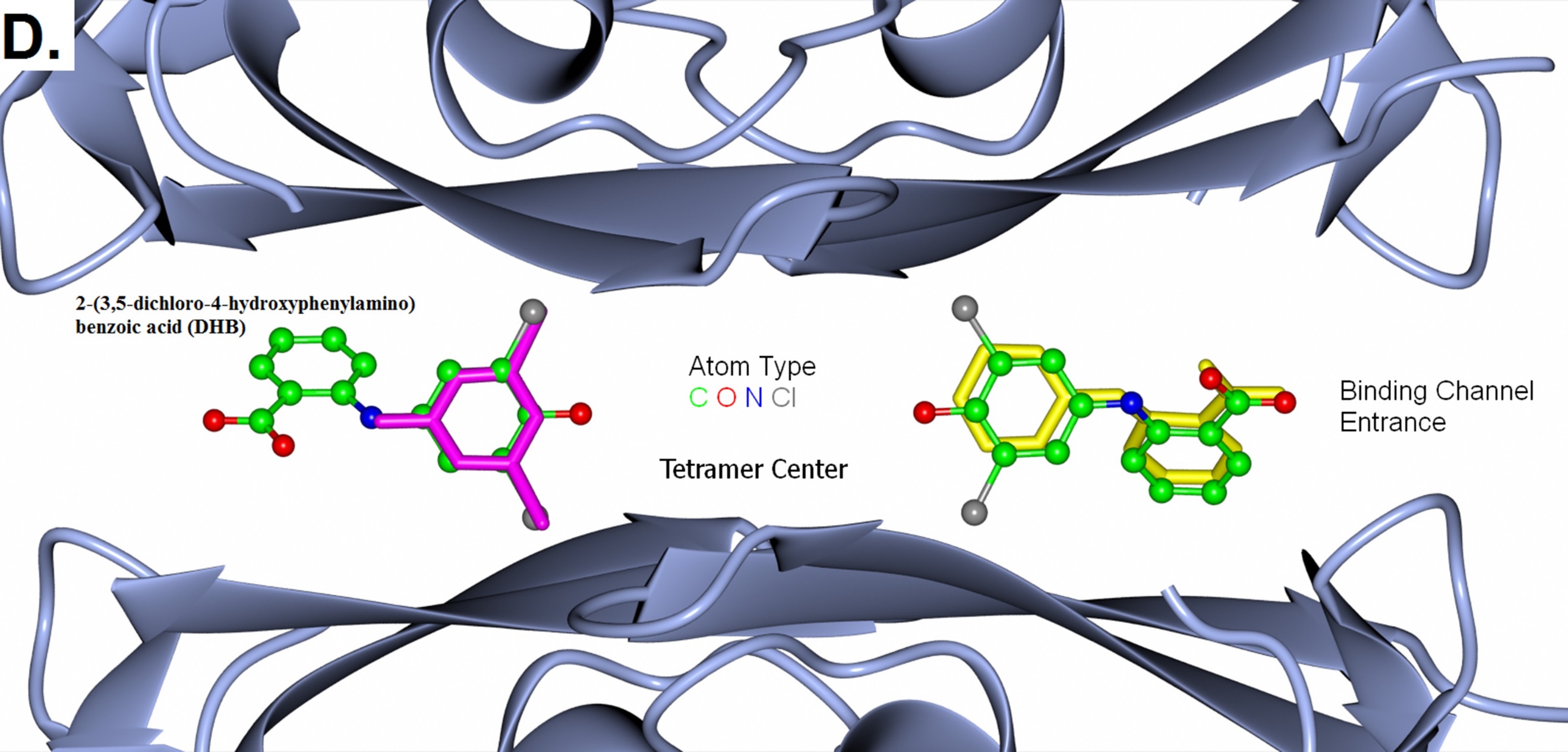

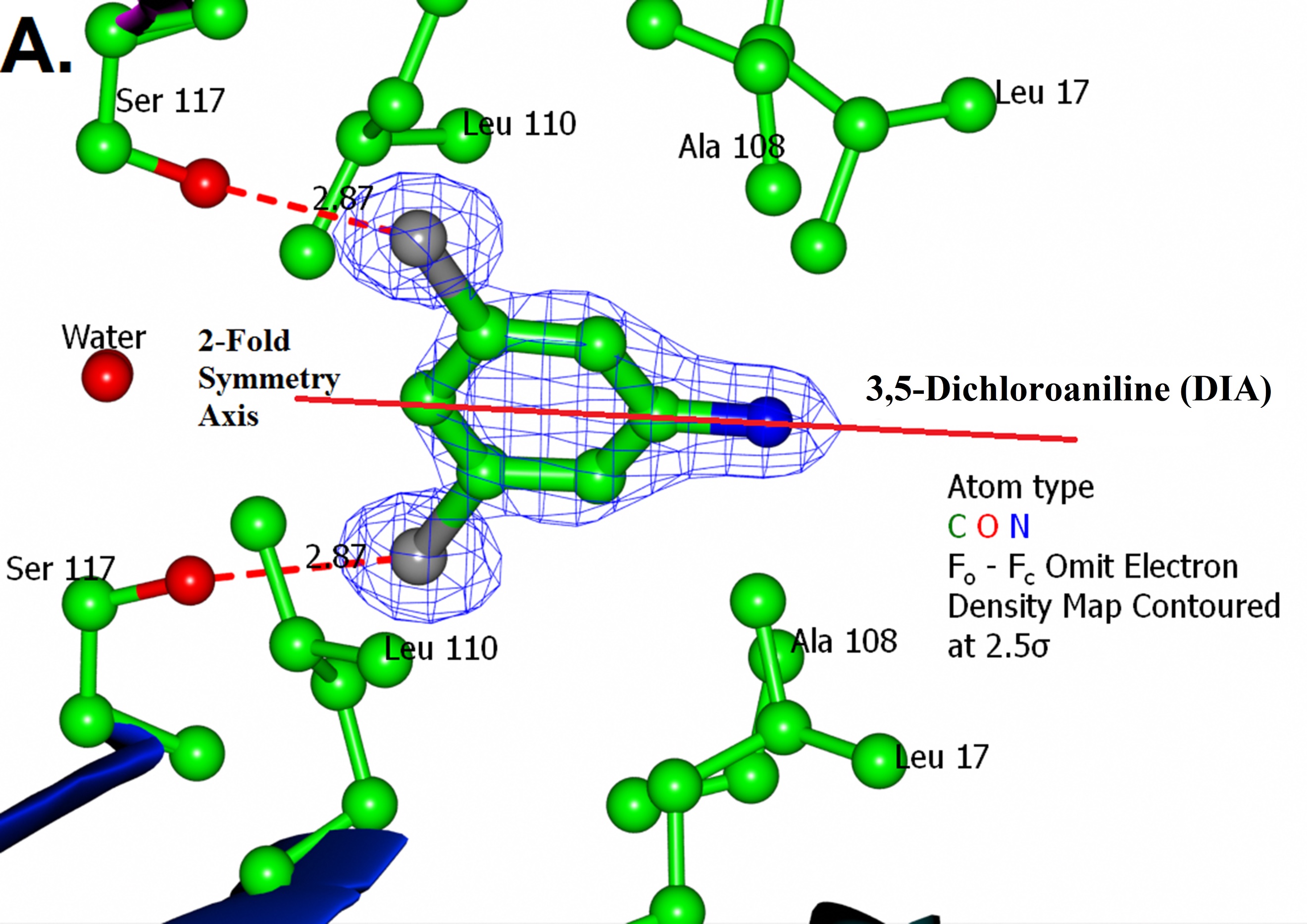

(**A.**) The chlorine atoms of 3,5-Dichloroaniline (DIA) are buried deep inside the binding channel and have formed halogen bonds to each of the Ser117 sidechains nearby. (**B.**) A chemical switch from amino to sulfonamide has caused the dichlorobenzene to face the opposite direction towards the binding channel entrance. (**C.**) Unlike DIA and DSF, N-Phenylanthranilic Acid (NPA) occupied both sites in the binding channel. Its diphenylamine is wrapped around by the sidechains of Leu110 and Leu17 whilst the carboxylate forms ionic interaction with Lys15. (**D.**) The crystal structures of DIA (magenta cylinders), NPA (yellow cylinders) and DHB (balls & sticks) in complex with TTR are superposed by Secondary Structure Matching. The corresponding atomic coordinates between these three compounds matched accurately. Numbers near dashed red lines are bond lengths in Angstroms.

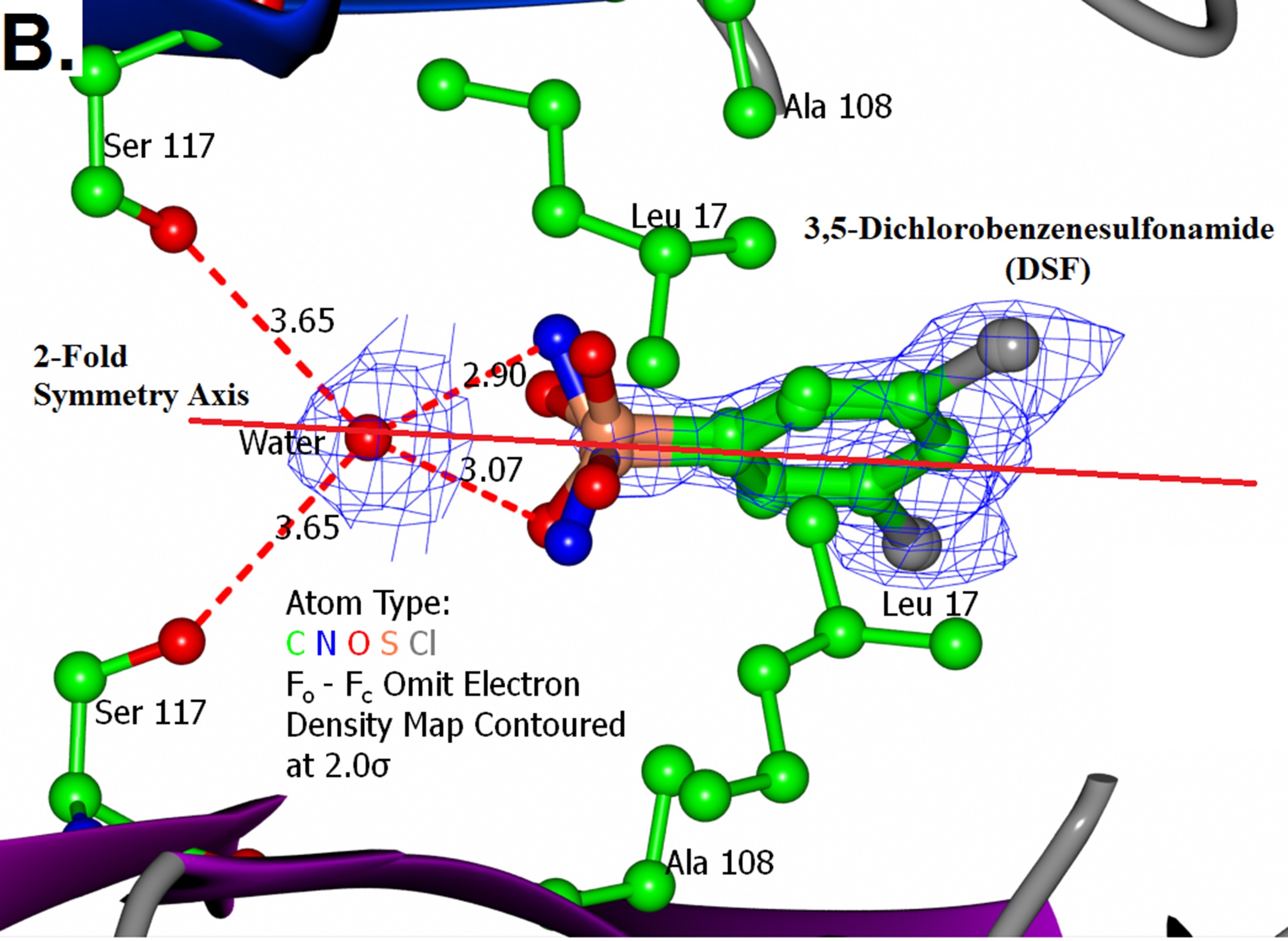
